# Type IV-A3 CRISPR-Cas systems drive inter-plasmid conflicts by acquiring spacers *in trans*

**DOI:** 10.1101/2023.06.23.546257

**Authors:** Fabienne Benz, Sarah Camara-Wilpert, Jakob Russel, Katharina G. Wandera, Rimvydė Čepaitė, Manuel Ares-Arroyo, José Vicente Gomes-Filho, Frank Englert, Johannes Kuehn, Silvana Gloor, Aline Cuénod, Mònica Aguilà-Sans, Lorrie Maccario, Adrian Egli, Lennart Randau, Patrick Pausch, Eduardo Rocha, Chase L. Beisel, Jonas S. Madsen, David Bikard, Alex R. Hall, Søren J Sørensen, Rafael Pinilla-Redondo

## Abstract

Type IV-A CRISPR-Cas systems are primarily encoded on plasmids and form multi-subunit ribonucleoprotein complexes with unknown biological functions. In contrast to other CRISPR-Cas types, they lack the archetypical CRISPR acquisition module and encode a DinG helicase instead of a nuclease component. Type IV-A3 systems are carried by large conjugative plasmids that often harbor multiple antibiotic-resistance genes. Although their CRISPR array contents suggest a role in inter-plasmid conflicts, this function and the underlying mechanisms have remained unexplored. Here, we demonstrate that a plasmid-encoded type IV-A3 CRISPR-Cas system co-opts the type I-E adaptation machinery from its clinical *Klebsiella pneumoniae* host to update its CRISPR array. Furthermore, we demonstrate that robust interference of conjugative plasmids and phages is elicited through CRISPR RNA-dependent transcriptional repression. By targeting plasmid core functions, type IV-A3 can prevent the uptake of incoming plasmids, limit their horizontal transfer, and destabilize co-residing plasmids, altogether supporting type IV-A3’s involvement in plasmid competition. Collectively, our findings shed light on the molecular mechanisms and ecological function of type IV-A3 systems and have broad implications for understanding and countering the spread of antibiotic resistance in clinically relevant strains.

## INTRODUCTION

CRISPR-Cas systems protect bacteria from invading mobile genetic elements (MGEs) by providing adaptive immunity. Essential to their memory acquisition is the conserved Cas1-Cas2 adaptation module, which excises short sequences (protospacers) adjacent to a motif (PAM) in invading MGEs and incorporates them into the CRISPR array as new spacers (**Fig. 1A**). Array transcription is followed by processing into mature CRISPR RNAs (crRNAs), which assemble with Cas proteins into crRNA-guided effector complexes that target complementary sequences, typically through nucleolytic activities ^1,2^.

**Figure 1.**
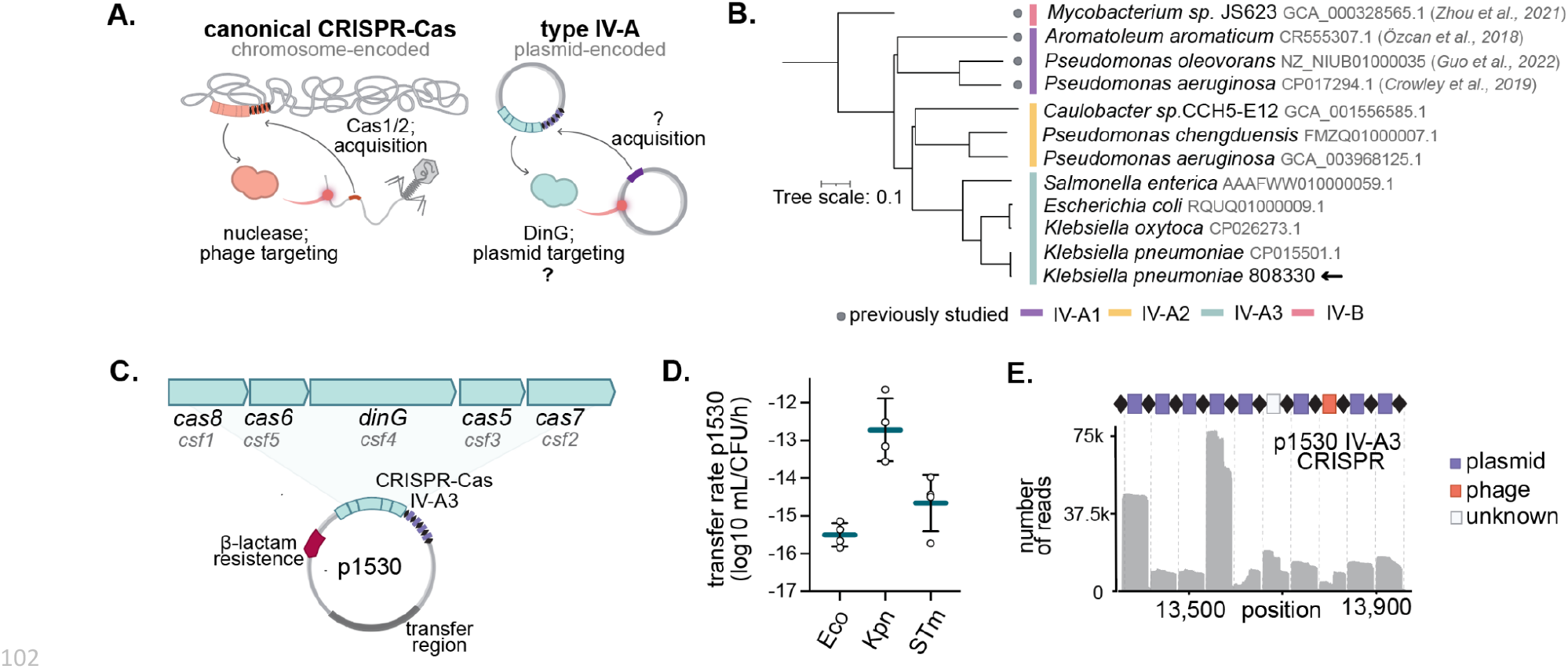
The *K. pneumoniae* type IV-A3 CRISPR-Cas system encoded on a conjugative ESBL-plasmid is functionally active. **A)** Comparison of canonical CRISPR-Cas systems (left) and type IV-A CRISPR-Cas (right). Most CRISPR-Cas systems are chromosomally encoded, contain Cas1/2 to acquire new spacers, and primarily target phages using associated nucleases. Type IV-A CRISPR-Cas are encoded on conjugative elements such as plasmids, lack Cas1/2 modules and carry a DinG helicase instead of a nuclease component. **B)** Phylogenetic tree showing the selected type IV-A3 from *K. pneumoniae* 808330 (arrow) among other type IV system representatives. Previously studied orthologs and taxonomic assignments of their hosts are indicated (gray dot). The phylogenetic tree was built using Csf2 (Cas7) protein alignments. **C)** Schematic of the type IV-A3 *cas* operon and CRISPR on the *K. pneumoniae* 808330 plasmid p1530. **D)** Mean rates of p1530 conjugation (blue lines) from its native host *K. pneumoniae* 808330 to different *Enterobacteriaceae* species: *E. coli* (Eco), *K. pneumoniae* (Kpn)*, and Salmonella enterica* Typhimurium (STm). The error bars indicate the SD (n=4). **E)** Schematic of the type IV-A3 CRISPR array carrying ten spacers (top), with their predicted origins indicated in purple (plasmid), orange (phage), or white (unknown). Small RNA-sequencing of the type IV-A3 CRISPR-Cas locus expressed in *E. coli* and mapped back to the p1530 CRISPR array (bottom).

Type IV CRISPR-Cas systems have remained largely understudied, in contrast to most other known CRISPR-Cas types ^3^. Like other Class 1 systems ^4^, they form multiprotein complexes ^5,6^ and are divided into distinct subtypes and variants based on their molecular architecture ^7,8^. Although type IV loci contain CRISPR arrays with varying spacer content, they lack Cas1-Cas2 adaptation modules, rendering their spacer acquisition mechanism enigmatic (**Fig. 1A**) ^7,9,10^. Type IV CRISPR-Cas systems also stand out for their consistent association with conjugative MGEs, such as plasmids and integrative conjugative elements (ICEs) ^5,7,9,11^, and in case of the type IV-A for featuring a 5’-3’ DNA helicase called DinG instead of an effector nuclease (**Fig. 1A**) ^3,7–9^.

Recent research has shed light on the potential mechanisms driving RNA-guided type IV CRISPR-Cas targeting. For example, the type IV-A CRISPR-Cas system (variant IV-A1) in *Pseudomonas oleovorans* has been shown to mediate DinG-dependent transcriptional repression of chromosomal targets ^12^. In contrast, type IV-A1 systems can facilitate the loss of a small vector plasmid even when the targeted region is outside an open reading frame ^10,12^, leaving open questions about the proposed CRISPR interference (CRISPRi) mechanism. Notably, type IV CRISPR arrays are enriched with spacers matching large conjugative plasmids, suggesting a unique role in inter-plasmid conflicts ^7,9,11^. However, their ecological role and whether and how they can interfere with conjugative plasmids remain unexplored.

Here, through a combination of molecular genetics, bioinformatics, and biochemical analyses, we functionally characterize a type IV-A3 CRISPR-Cas system encoded on a *Klebsiella pneumoniae* conjugative plasmid. Our results reveal that type IV-A3 can acquire spacers by co-opting the host-derived type I-E adaptation machinery. Additionally, we show that crRNA-guided targeting can mediate the loss of conjugative plasmids through transcriptional repression of plasmid core functions and demonstrate that this silencing activity can be repurposed to re-sensitize bacteria to antibiotics. Because type IV-A3 systems are widespread among the pervasive and opportunistically pathogenic *K. pneumoniae ^7,11,13^*, our findings have important implications for understanding plasmid-driven adaptation, including prevention and dissemination of antibiotic resistance and virulence factors.

## RESULTS

### A clinical *K. pneumoniae* conjugative plasmid encodes an active type IV-A3 CRISPR-Cas

Previous research and our analyses indicate that type IV-A3 systems are carried by large conjugative plasmids in *Enterobacteriaceae* (median size 280 kb), predominantly within the *Klebsiella* genus (91 %; **Suppl. Fig. S1A-D**) ^7,11^. These plasmids are usually cointegrates of IncHI1B/IncFIB replicons (53 %; **Supp. Fig. S1E**) and frequently carry one or more antibiotic resistance genes (58 %, on average 11 resistance genes/plasmid; **Supp. Fig. S1F-H**) ^13^.

To investigate the biological function and molecular mechanisms driving adaptation and interference in type IV-A CRISPR-Cas, we aimed to establish a model system with ecological and clinical relevance. We selected the clinical isolate *K. pneumoniae* 808330 (sequence type, ST182), which has a chromosomal type I-E CRISPR-Cas system and harbors plasmid p1530 that encodes a type IV-A3 CRISPR-Cas system (**Fig. 1B-C, Suppl. Fig. S2**). The IncHI1B/IncFIB p1530 is 205 kb and encodes the extended-spectrum β-lactamase (ESBL)-gene *bla*_CTX-M-15_ that has spread globally and confers resistance to important 3^rd^ generation cephalosporins ^14,15^. Through conjugation experiments, we confirmed the ability of p1530 to transfer efficiently from its natural host into another *K. pneumoniae* strain and also other clinically relevant *Enterobacteriaceae* species (**Fig. 1D**).

Our analysis of the CRISPR spacer contents in p1530 confirmed the strong preference for targeting other conjugative plasmids predicted across type IV CRISPR-Cas systems (**Fig 1E, Suppl. Fig. S2, Suppl. Table S1**) ^7^. Small RNA sequencing of *E. coli* expressing the type IV-A3 from a plasmid showed the production of mature crRNAs (**Fig. 1E, Suppl. Fig. S3A**). We then confirmed the constitutive expression and crRNA maturation of the type IV-A3 and type I-E CRISPR-Cas systems in their native *Klebsiella* host through total and small RNA sequencing, respectively (**Suppl. Fig. S3B-C**). Finally, heterologous protein expression and purification revealed the formation of a ribonucleoprotein complex, containing the Cas proteins Cas8 (Csf1), Cas6 (Csf5), Cas5 (Csf3), and multiple Cas7 (Csf2), but lacking DinG (Csf4) at a stoichiometry indicating co-precipitation (**Suppl. Fig. S4**). This shows that the type IV-A3 Cas proteins assemble into a multisubunit complex and indicates that DinG may be recruited to the interference complex *in trans* after target binding, similar to what has been observed for Cas3 in CRISPR-Cas type I^4^. These results collectively suggest that the *K. pneumoniae* type IV-A3 system in p1530 is both functional and suitable as a model system.

### *In trans* use of Cas1/2e facilitates spacer acquisition in type IV-A3 CRISPR arrays

Type IV-A loci lack adaptation modules despite their association with CRISPR arrays with varying spacer content, prompting questions about the spacer acquisition mechanism (**Fig. 1A**). Notably, type IV-A3 systems are frequently found in strains that encode chromosomal type I-E systems, and our previous bioinformatic analyses revealed significant similarities in their CRISPR repeats and leader sequences ^7^. To investigate the potential functional interplay between type IV-A3 CRISPR arrays with type I-E adaptation modules, we expressed *K. pneumoniae* Cas1e and Cas2e (Cas1/2e) in *E. coli* harboring the type IV-A3-encoding plasmid p1530. To enhance rare spacer acquisition events, we electroporated cells with 35 bp double-stranded DNA oligos as protospacers (PS) containing the canonical type I-E 5’-AAG-3’ spacer acquisition motif (SAM) (**Fig. 2A, Suppl. Fig. S5A**) ^16^. PCR analysis revealed Cas1/2e-dependent array expansion (**Fig. 2B, Suppl. Fig S5B**), and Sanger sequencing confirmed integration of the electroporated protospacer into the leader-repeat junction of the type IV-A3 CRISPR array in p1530 (**Fig. 2B)**. Notably, the 3’-guanine of the SAM was incorporated into the CRISPR array together with the protospacer sequence (**Fig. 2B**), which is a distinctive characteristic of type I-E adaptation ^17–19^.

**Figure 2:**
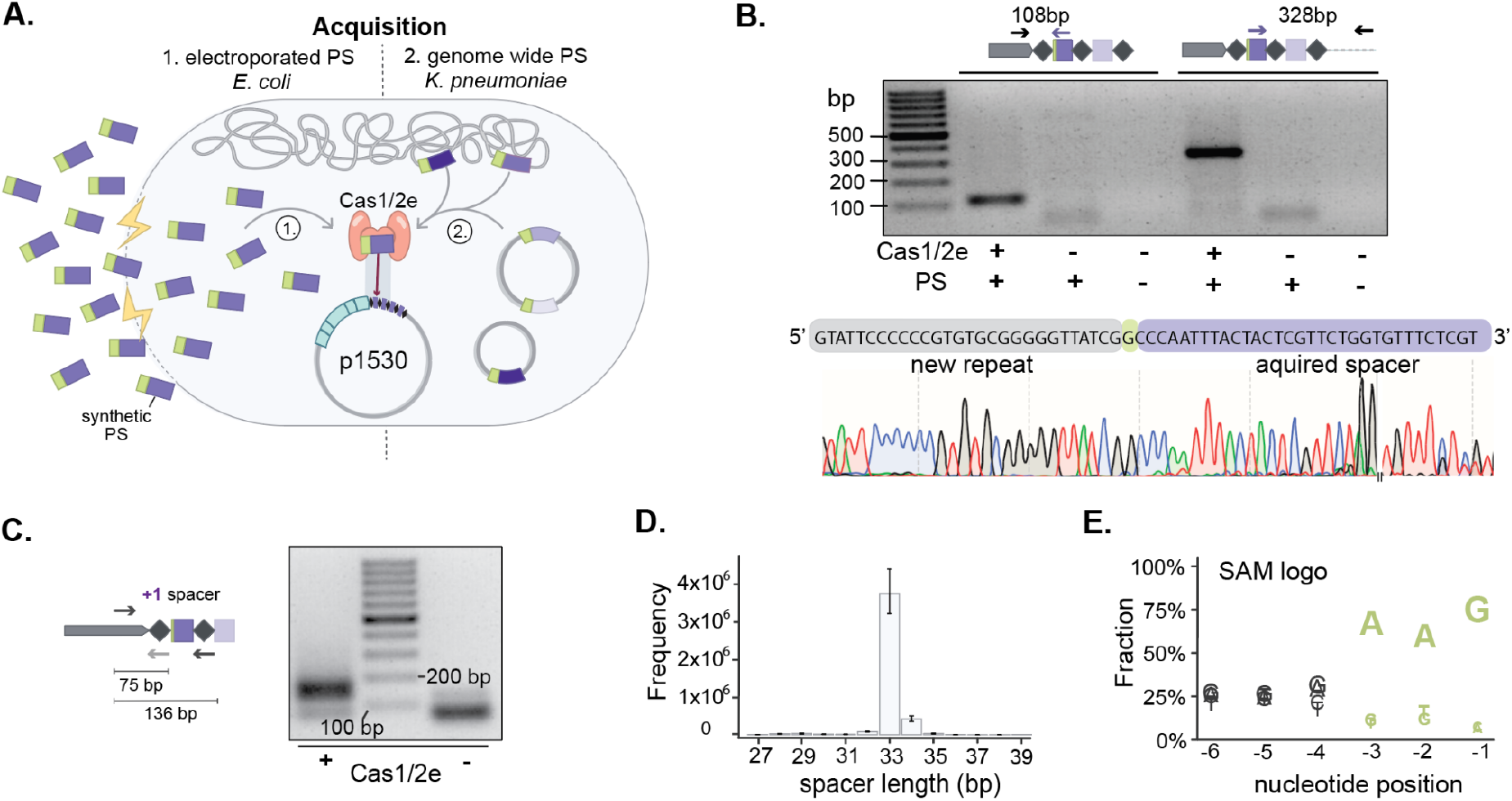
Type I-E adaptation machinery facilitates spacer acquisition in type IV-A3 CRISPR loci. **A)** Schematic of the two acquisition experiment setups; experiment 1: *E. coli* harboring the p1530-encoded type IV-A3 CRISPR-Cas system and electroporated with double-stranded DNA protospacers (PS, purple) with a 5’-AAG-3’ SAM (green). Experiment 2: *K. pneumoniae* with a targeting-deficient p1530-encoded type IV-A3 system (Δ*dinG*). In both experiments, the Cas1/2e adaptation module was expressed from a plasmid. **B)** Experiment 1: Amplicons of leader proximal CRISPR array region and corresponding Sanger-sequencing results (bottom). The schematics, gel, and Sanger results show the amplification of the CRISPR array upstream and downstream of the newly acquired spacer. The break in the Sanger sequence trace indicates the assembly of sequences from opposing directions. **C)** Experiment 2: Detection by PCR of Cas1/2e dependent genome-wide spacer acquisition in *K. pneumoniae* by amplicon deep-sequencing. The gel and schematics show amplification during the second PCR of CAPTURE of the leader-proximal end, 136 bp for elongated arrays (left) and 75 bp for the leader amplification (right). Black arrows indicate primer annealing sites and the gray arrow indicates a secondary binding site. **D)** Mean number of integrated spacers by length (x-axis) as bars with error bars indicating the 95 % confidence interval (n=3). **E)** Sequence logo of the SAM as determined by the genome-wide spacer acquisition assay. Nucleotide abundance is shown as the mean fraction (n=3) at positions -6 to -1 of the acquired 33 bp-spacers.

To further characterize Cas1/2e-mediated spacer acquisition into type IV-A3 arrays and the corresponding SAM, we overexpressed Cas1/2e in *K. pneumoniae* 808330 harboring p1530 with a targeting-deficient type IV-A3 (Δ*dinG*) to allow for genome-wide spacer acquisition (**Fig. 2A**). We PCR-amplified and deep-sequenced expanded CRISPR arrays (**Fig. 2C, Suppl. Fig. S5C**) ^20^ and mapped the acquired spacers back to the *K. pneumoniae* 808330 genome. Consistent with the acquisition in I-E CRISPR-Cas systems, the majority of acquired spacers were 33 bp in length (85 %, n= 13M total; **Fig. 2D**) and originated from genomic positions next to a 5’-NAAG-3’ SAM (position -3 to -1, 49 %, total percentage across all spacers; **Fig. 2E**) ^17–19^. Furthermore, we observed a preference for the acquisition of spacers from plasmids in the cell (**Suppl. Fig. S5D**), consistent with previous reports ^21^, and no preference for the coding and template strands (**Suppl. Fig. S5D-E**). Together, our findings demonstrate that plasmid-encoded type IV-A3 CRISPR-Cas systems can use host-derived Cas1/2e to acquire new spacers.

### IV-A3 targeting interferes with horizontal transfer and stability of conjugative plasmids

Type IV-A3 CRISPR-Cas spacers exhibit a prominent bias towards targeting other large conjugative plasmids (median size 136 kb, 67 % conjugative; **Suppl. Fig. 6A-D**), leading to speculation about their biological role in mediating inter-plasmid conflicts ^7,9,11,22^. The targeted plasmids, which are predominantly found in *Klebsiella* and span various Inc groups, are often replicon cointegrates such as IncFII/IncFIB (**Suppl. Fig. 6E-G**). Of particular importance, they commonly harbor multiple antimicrobial resistance genes (57%; **Suppl. Fig. S6H-I**).

We reasoned that plasmid competition mechanisms may act on the horizontal or vertical inheritance of other plasmids, compromising their long-term stability in bacterial populations. To test whether type IV-A3 can mediate inter-plasmid competition, we leveraged three experimental setups to explore the impact of type IV-A3 targeting on pKJK5, a broad-host-range IncP-1 antibiotic-resistance plasmid (**Fig. 3A-E**) ^23,24^. We designed crRNAs targeting both DNA strands of pKJK5 in selected regions involved in plasmid replication (replication initiation, *trfA*), inheritance (partitioning, *parA*), conjugation (transfer initiation, relaxase *traI*, and origin of transfer, *oriT*), and expression of a heterologous green fluorescent protein (*gfpmut3*, hereafter *gfp*) (**Fig. 3A**, **Suppl. Fig. S7A-C**).

**Figure 3.**
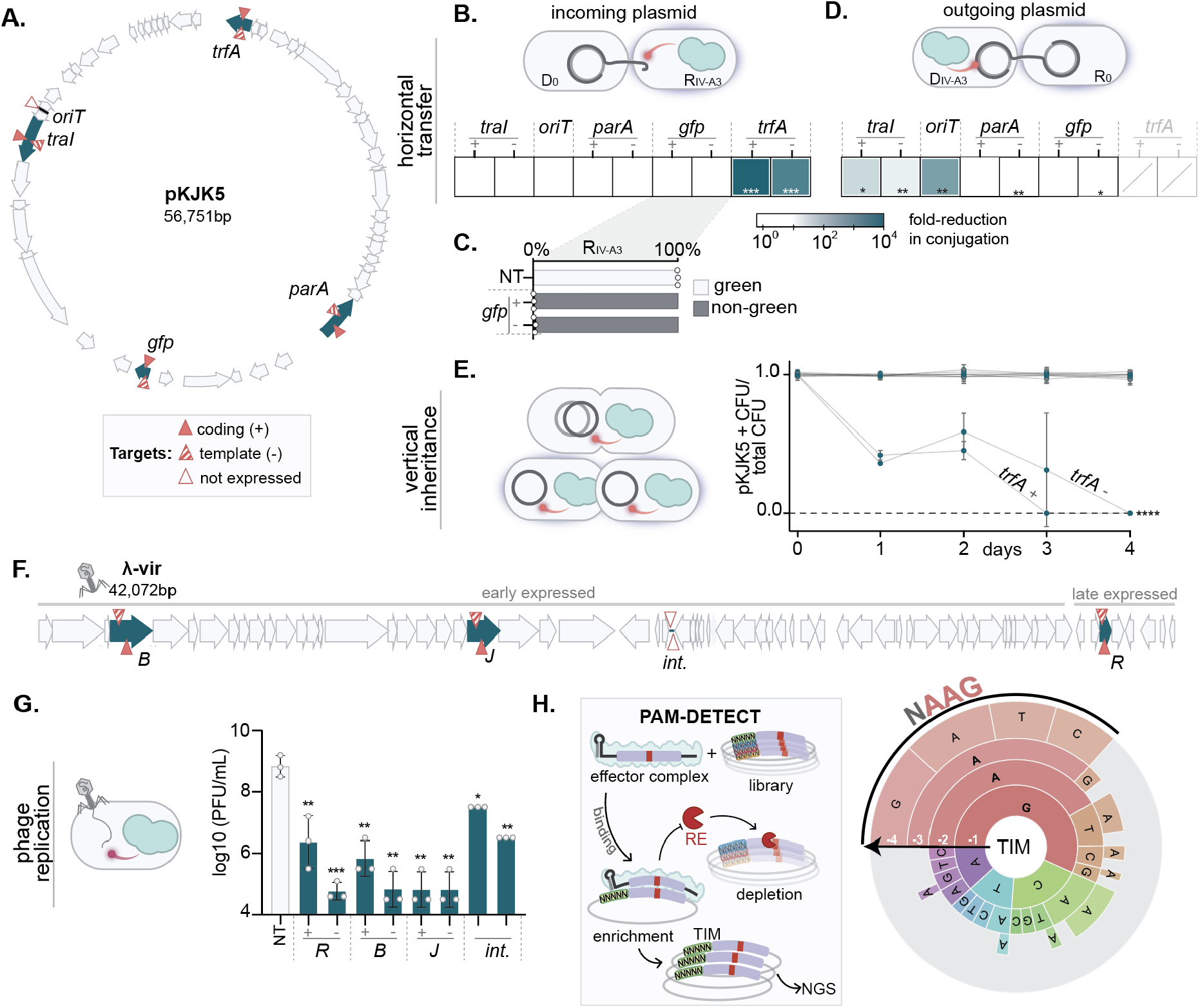
IV-A3 mediates crRNA-directed interference with conjugative plasmids and phages. **A)** Gene map of pKJK5 indicating the regions targeted by type IV-A3 in blue. Red triangles represent the approximate location of protospacers. **B)** Type IV-A3 interference in *E. coli* recipients R_IV-A3_. **C)** Evaluation of GFP fluorescence under type IV-A3 targeting in R_IV-A3_. Bars represent the mean of transconjugants emitting green and non-green signals, comparing a non-targeting (NT) crRNA control to crRNAs targeting *gfp.* **D)** Type IV-A3 interference in *E. coli* donors D_IV-A3_. Evaluation of *trfA* targeting was not feasible due to the instability of pKJK5 while expressing type IV-A3 CRISPR-Cas (crossed, gray-shaded squares). The conjugation efficiency of pKJK5 in **B** and **D** is shown as the conjugation reduction compared to the NT control. **E)** Plasmid maintenance assay showing pKJK5 stability under type IV-A3 targeting in the absence of pKJK5-selection over ∼40 generations. The dotted line indicates the detection limit of the assay. Blue dots show the mean of four biological replicates with error bars as SD (n=4). **F)** Genome map of λ-vir indicating the regions targeted by type IV-A3 in blue. Target sites are shown as in **A**. Early and late expressed regions are indicated. **G)** Type IV-A3 interference of λ-vir infection in *E. coli* determined as plaque forming units (PFU)/ mL. Bars show the mean values and error bars indicate the SD. **H**) Schematic of PAM-DETECT method on the left. TIM wheel on the right shows the mean sequence motif recognized by type IV-A3 interference complex in 5’ > 3’ from outer to inner position (n=2). The size of the arc for each nucleotide position corresponds to its relative enrichment within the TIM library. Individual sequences comprising at least 2 % of the PAM wheel are shown. P values in panels **B** and **D-F** represent two-sample Student’s t-Tests of log10 transformed T/(R+T) for panels **B** and **D**, pKJK5^+^ fraction for **E** and PFU/mL for **G**, comparing each targeting treatment to the NT control (n=3 if not stated otherwise). **** P ≤ 0.0001; *** P ≤ 0.001; ** P ≤ 0.01; *P ≤ 0.05.

We first asked whether plasmid targeting could limit pKJK5 establishment in a type IV-A3-expressing recipient strain (RIV-A3) upon conjugation from a donor strain (D_0_) (**Fig. 3B**). When targeting the *trfA* gene, which is essential for plasmid replication, conjugation efficiency was reduced by over four orders of magnitude while targeting the non-essential sites had no effect on pKJK5 establishment (**Fig. 3B, Suppl. Fig. S7D**). Notably, >95 % of transconjugants with targeted *gfp* did not emit green fluorescence, supporting the interference mechanism by transcriptional repression proposed for type IV-A1 (**Fig. 3C**)^12^. We then investigated whether targeting pKJK5 in type IV-A3-expressing donors (DIV-A3) could hinder its transfer to a recipient strain (R_0_) (**Fig. 3D**). Targeting the *oriT* and the *traI* gene, which are required for conjugation, substantially reduced horizontal plasmid transfer (**Fig. 3D, Suppl. Fig. S7E**). This is in contrast to the plasmid-incoming experiment, where targeting the same regions did not affect conjugation efficiencies (**Fig. 3B, Suppl. Fig. S7D**). Finally, we tested whether type IV-A3 interference could destabilize pKJK5 in a plasmid stability assay (4 days; ∼40 generations) in the absence of pKJK5-specific selection. In this assay, pKJK5 was only lost when targeting the *trfA* gene, confirming its essentiality (**Fig. 3E, Suppl. Fig. S8A**). We further explored whether type IV-A3 targeting incurred a growth disadvantage on cells with a targeted plasmid, as shown for other anti-plasmid systems that function through abortive infection ^25,26^. However, there was no qualitative difference in population growth upon targeting (**Suppl. Fig. S8B**). Our experiments demonstrate that type IV-A3 CRISPR-Cas can effectively limit both the transfer and stability of natural conjugative plasmids in bacterial populations, regardless of the targeted strand, and are consistent with a natural CRISPRi mechanism.

### IV-A3 interferes with phage propagation

A small fraction of spacers in type IV CRISPR arrays are predicted to match phage sequences (**Fig. 1E**, **Suppl. Fig. S2, S6A, Suppl. Table S1**) ^7,9,10^, suggesting a selective advantage for plasmids to retain these spacers. To evaluate this, we challenged *E. coli* with phage λ-vir and designed type IV-A3 crRNAs against both DNA strands of the λ-vir genome at four selected positions, including early and late expressed genes, and an intergenic region (**Fig. 3F**). Type IV-A3 interference reduced the ability of λ-vir to propagate in its host for up to five orders of magnitude (**Fig. 3G**). Interestingly, interference with phage infection was significant but less pronounced when targeting the intergenic region (**Fig. 3G**). These results indicate that type IV-A3 CRISPR-Cas systems can robustly target phages and suggest that type IV-A3-carrying plasmids can enhance their own fitness by protecting their hosts from phage predation.

### IV-A3 interference requires the presence of a target interference motif

Mutational evasion of CRISPR-targeting by phage evasion has provided valuable insights into the mechanistic constraints of CRISPR-Cas systems, revealing that the PAM and seed (PAM proximal region, important for target identification initiation) in the protospacer are essential for interference ^27–29^. To deepen our mechanistic understanding of type IV-A3 targeting, we isolated and analyzed a set of λ-vir variants capable of escaping interference. We found λ-vir evaded targeting by mutations in the 2^nd^ and 3^rd^ positions of the 5’-AAG-3’ PAM, in the seed, or by deleting the seed and PAM region (**Suppl. Fig. S9A-D**), altogether suggesting a reliance on the stringent recognition of a target interference motif (TIM).

To further characterize the TIM requirements of type IV-A3, we used an *in vitro* cell-free transcription-translation (TXTL) assay for PAM determination (PAM-DETECT) based on the restriction enzyme-dependent depletion of protospacer sequences without a recognized TIM ^30^. This revealed a pronounced dependence of the interference complex on the recognition of a 5’-NAAG-3’ TIM for protospacer binding **(Fig. 3H)**. The identified TIM exhibits striking consistency with the 5’-AAG-3’ SAM determined in our Cas1/2e dependent acquisition experiments (**Fig. 3H**, **Fig. 2E**), highlighting the compatibility in functional requirements between the type I-E adaptation machinery and type IV-A3 interference complex. Finally, we found that λ-vir also overcame interference through deletion of the region encoding the targeted lysozyme gene *R,* and subsequently acquiring a functional homolog present in other coliphages (**Suppl. Fig. S9E**). Together, these results highlight the strong selective pressure exerted by type IV-A3 interference with λ-vir and the stringent recognition of a TIM for effective targeting.

### DinG is essential for blocking expression once transcription has initiated

The above results and a recent study suggest that type IV-A systems elicit target interference through transcriptional repression ^12^, shedding light on the targeting mechanism. However, the mechanistic role of the associated DinG helicase remains unclear despite its suggested ATP-dependent 5’-3’ DNA helicase activity on the target ^8^ and its proposed essentiality in this process ^10,12^. To shed light on the relevance of DinG during interference, we used three variants of the type IV-A3 CRISPR-Cas system: wildtype, a DinG knockout mutant (Δ*dinG*), and a catalytically inactive helicase mutant (D215A/E216A, *dinGmut;* **Fig. 4A**) ^10^.

**Figure 4:**
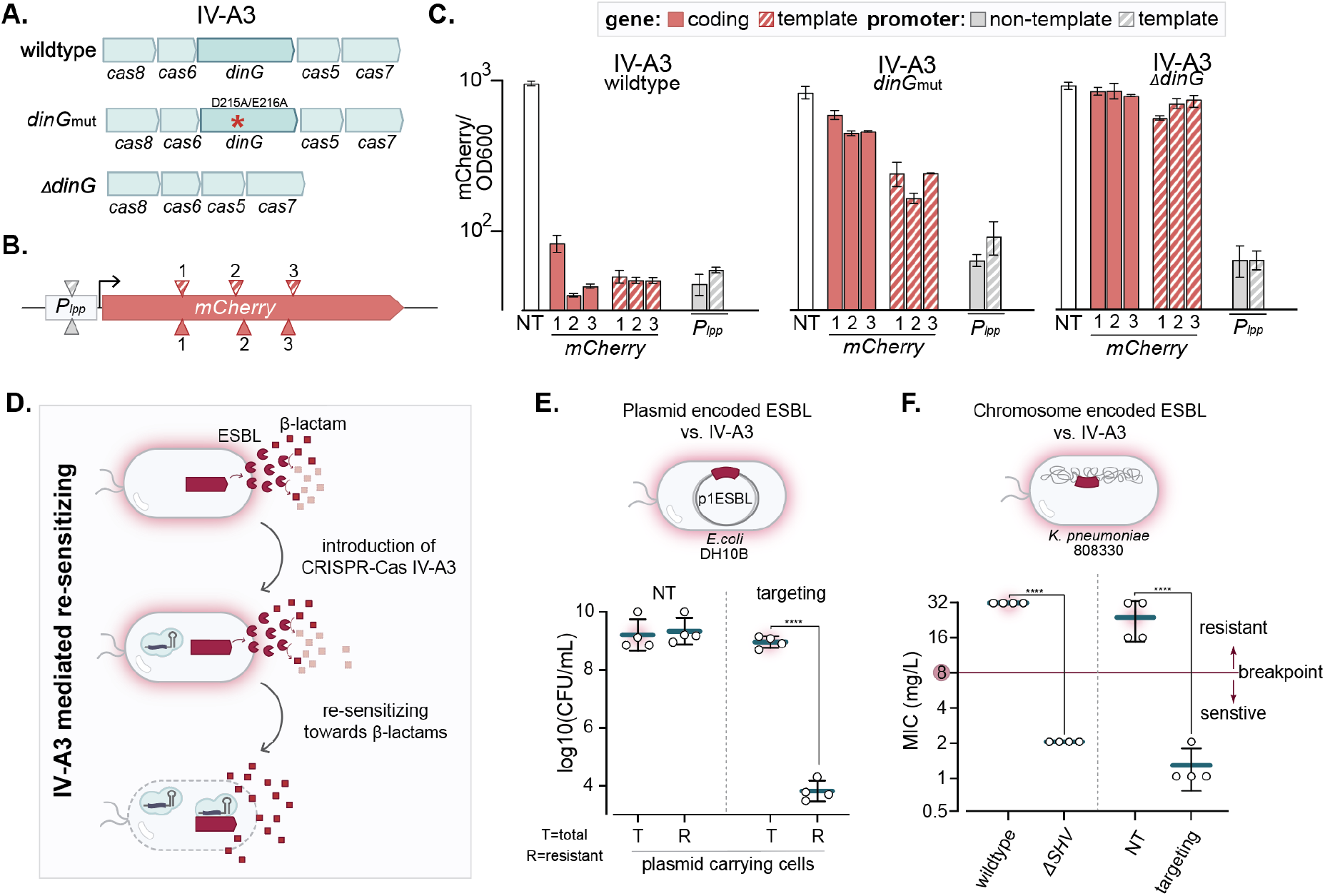
IV-A3 interference functions through transcriptional repression and can re-sensitize antibiotic-resistant bacteria. **A)** Schematic of the three type IV-A3 CRISPR-Cas variants used in the *mCherry* reporter targeting assay; wildtype, *dinGmut* with a catalytically inactive DinG, and the Δ*dinG* knockout mutant. **B)** Illustration of the mCherry reporter construct. Triangles represent the approximate location of protospacers within *mCherry* (red) and the *Plpp* promoter (gray), where the crRNA hybridizes to the coding (full) and template (crosshatched) strands **C)** *In vivo* transcriptional repression assay. The mean mCherry fluorescence signal is normalized to bacterial OD_600_ (y-axis) and the targeted positions are shown on the x-axis. Error bars indicate SD. A linear model indicated that IV-A3 variant, target position (promoter vs. gene), and strand significantly contribute to relative mCherry levels (*R^2^* = 0.98, P < 0.0001 for all factors) and a bidirectional stepwise linear regression that target position is the most important predictor, followed by IV-A3 variant, and then strand (n = 3). **D)** Diagram illustrating the process of restoring β-lactam sensitivity in ESBL-producing strains through crRNA-guided type IV-A3 CRISPR-Cas gene silencing. **E)** Reversal of ESBL-mediated antibiotic resistance encoded on p1ESBL carried by *E. coli* DH10B shown as the log_10_ of colony forming units (CFU)/ mL under type IV-A3 interference (targeting) and the NT control. Lines represent the mean of the plasmid-carrying cells: total population of plasmid-carrying cells (T) and the β-lactam resistant subpopulation thereof (R) (n= 4). Error bars indicate the SD. **F)** MIC of Amp for *K. pneumoniae* 808330 under type IV-A3 mediated transcriptional repression of *bla*_SHV-187_ (right) and for a Δ*SVH* mutant (left). The EUCAST susceptibility breakpoint (8 mg/L) is indicated in red. Datapoints shown at 32 mg/L are ≥ 32 mg/L and error bars indicate SD. The p-value in panels **E** and **F** represents a two-sample Student’s t-Tests, of the log_10_ transformed CFU/mL for E and MIC values for F (n = 4 for both). **** P ≤ 0.0001.

We then assessed the ability of the variant systems to target fluorescent reporter genes at various intragenic positions and promoter sites, using an *in vivo* setup (chromosomal *mCherry*; **Fig. 4B**) and an *in vitro* TXTL setup (*degfp*; **Suppl. Fig. S10A-B**). Using the wildtype IV-A3 CRISPR-Cas system, targeting within the open reading frame of the reporter genes consistently resulted in a robust decrease of fluorescent signal (**Fig. 4C**; **Suppl. Fig. S10C**). In contrast, interference was completely abolished in the absence of DinG (Δ*dinG*), supporting its requirement for gene silencing. With the *dinGmut* variant, we found a reduced mCherry signal when crRNAs hybridized to the template strand but not when crRNAs hybridized to the coding strand *in vivo*. Targeting in the promoter region, however, always resulted in a strong reduction of reporter signal, independent of the DinG variant (**Fig. 4C, Suppl. Fig. S10C**). Indeed, we found that purified type IV-A3 complexes bound strongly (at a low nM apparent dissociation constant K_D_) to a cognate double-stranded DNA target in the absence of DinG (**Suppl. Fig. S11A-C**). This shows that initial DNA target binding is independent of DinG and suggests that the reduced reporter signal, observed upon targeting the promoter region of mCherry (**Fig. 4C**), arises from blocked transcription initiation upon type IV-A3 ribonucleoprotein complex binding. Together, these results suggest that the type IV-A3 complex without DinG is sufficient to prevent transcription initiation, while DinG is crucial to mediate gene repression in transcribed regions.

### IV-A3 mediated re-sensitization of antibiotic-resistant bacteria

Due to the growing spread of antimicrobial resistance in pathogenic strains and its impact on global health ^31^, there is an urgent need to develop alternative strategies such as restoring antimicrobial susceptibility ^32–34^. Our findings highlight that type IV-A3 shows promise as a programmable tool for transcriptional repression, which is particularly noteworthy given the natural propensity of type IV-A3 systems to target conjugative multidrug-resistance plasmids carried by clinical pathogens (**Suppl. Fig. S6F, H-I**).

To investigate the suitability of type IV-A3 for re-sensitizing bacterial strains to antimicrobials, we guided the effector complex towards β-lactam resistance genes (**Fig. 4D**). By targeting the ESBL-gene (*bla*_CTX-M15_) encoded on the clinical *E. coli* plasmid p1ESBL (**Fig. 4E**) ^35,36^, we restored the strain’s susceptibility to the extended-spectrum β-lactam antibiotic ampicillin (Amp). Importantly, the targeted plasmid was maintained in re-sensitized cells, consistent with the transcriptional repression mechanism (**Fig. 4E**). With a broth microdilution minimum inhibitory concentration (MIC) assay, we further demonstrated that targeting the chromosomal *bla*_SHV-187_ in *K. pneumoniae* 808330 reduced the MIC value below the EUCAST clinical susceptible breakpoint (8 mg/L for Amp, v13.0, 2023-01-01; **Fig. 4F**). The resistance reduction was similar to that caused by the SHV-187 null-mutation (Δ*SVH*) (**Fig. 4F**) or the addition of the β-lactamase inhibitor clavulanic acid in a disc diffusion assay (**Suppl. Fig. S12A-B**). Together, our results underscore the programmability of type IV-A3 systems for silencing target genes of interest and exemplify their use for combating antimicrobial resistance.

## DISCUSSION

CRISPR-Cas systems encoded on MGEs frequently lack adaptation modules ^22,37–39^ and how they acquire new spacer memory has remained enigmatic. Our experiments support a model in which type IV-A3 CRISPR-Cas systems can overcome this limitation by employing host-derived I-E Cas1/2e proteins (**Suppl. Fig. S13**). Since the adaptation machinery has a strong preference for sampling MGEs ^40^, type IV systems may reduce plasmid self-targeting costs while enabling acquisition occasionally, when compatible adaptation modules become available *in trans*. Our findings reveal a striking functional overlap in PAM recognition preferences between the type I-E adaptation and type IV-A3 interference complexes (SAM and TIM, respectively), underscoring remarkable co-evolution of DNA motif specificity among distinct CRISPR-Cas types. We speculate that other mobile CRISPR-Cas systems, including CRISPR-associated transposons ^38,41–43^ and certain phage- ^39,44,45^ and plasmid-encoded loci ^22^, may similarly co-opt host adaptation machinery to acquire spacer content. A growing body of work is revealing the frequent carriage of diverse anti-phage defense systems by MGEs ^46,47^, suggesting that such functional complementarity with chromosomal loci may be a widespread phenomenon beyond CRISPR-Cas.

Despite growing evidence that type IV-A CRISPR-Cas systems are primarily involved in inter-plasmid conflicts ^7–9^, this hypothesis has remained unexplored. Our findings demonstrate that type IV-A3 can effectively block horizontal transfer and vertical inheritance of conjugative plasmids in bacterial populations by silencing essential plasmid functions. The benefits of licensing interference through non-nucleolytic activities are unclear. However, in contrast to nucleolytic CRISPR-Cas systems, transcriptional repression may be less likely to trigger DNA-damage-induced SOS response that impairs host growth ^48–50^ and plasmid fitness ^51^. This advantage may extend to other MGE-encoded CRISPR-Cas systems lacking nuclease activity, such as types V-M ^45^ and V-C ^52^, the helicase-associated type I-C variant ^39^, as well as other type IV systems ^7^.

Non-nucleolytic interference may present further advantages, including the capacity to acquire spacers that target chromosomal genes without causing toxic effects. Furthermore, this mechanism could allow plasmids to selectively retain spacers that manipulate the host’s or other co-residing MGE’s transcriptional profiles to their advantage. In support of this, Guo et al. 2022 reported the repression of the chromosomal pilus biogenesis gene PilN by a plasmid-encoded type IV-A1 system. Interestingly, a significant proportion of type IV CRISPR spacers match plasmid conjugation genes ^7^, which are also involved in pilus formation. For example, phages using pili as receptors are widespread ^53^, and it is possible that type IV-driven pili repression enhances plasmid fitness by preventing phage entry into host cells.

Albeit dependent on the presence and catalytic integrity of DinG, we demonstrate that type IV-A3 can robustly interfere with transcription elongation when targeting both the coding and template strands (**Suppl. Fig. S13**). This finding highlights the potential of type IV-A3 as a strand-independent CRISPRi tool that contrasts the conventional nuclease-deficient Cas9 (dCas9) ^54,55^, dCas12 ^56,57^ and Cascade ^58^, which are mostly restricted to the coding strand. As a proof-of-concept demonstration, we show that type IV-A3 gene silencing can be repurposed to re-sensitize bacteria to antibiotics, including high-risk clinical *K. pneumoniae* strains resistant to last resort β-lactams ^59^. We further showcase that the targeting of plasmid-encoded accessory genes does not cause plasmid loss, highlighting the distinctive potential of type IV-A3 to prevent the emergence of CRISPR-Cas inactivating mutations ^60,61^. Indeed, such unwanted mutations are particularly favored when targeting natural plasmids for removal, as they frequently encode addiction systems that select for their maintenance in the population ^62,63^. We anticipate that further investigations of the molecular mechanisms underlying type IV CRISPR-Cas systems will present further opportunities for harnessing their unique crRNA-guided properties in biotechnological applications.

## MATERIALS AND METHODS

### Bacterial strains, phages, and growth conditions

Bacterial strains phages, and plasmids used in this study are listed in **Table S3 and 4**. We screened the *Klebsiella spp.* collection from the University Hospital Basel, Switzerland to identify *K. pneumoniae* 808330, which was isolated in 2017 from a rectal swab taken during a hospital hygiene screening. We performed library preparation for *K. pneumoniae* 808330 with a Nextera XT Kit to sequence on an Illumina NextSeq500 (paired end, 2 × 250 bp) and a rapid barcoding sequencing kit (SQK-RBK004). We then proceeded to sequence with an Oxford Nanopore MinION system (FLO-MIN-106 flow cell). To generate a hybrid assembly, we used Unicycler v0.4.8 ^64^ and annotated CRISPR-Cas systems using CRISPRCasTyper ^65^, plasmids with Plasmidfinder and MOB-suite ^66,67^, and resistance genes by blasting against the CARD database ^68^. Unless stated otherwise, bacterial cultures grew at 37 °C and under agitation (180 rpm) in lysogenic broth (LB) medium, supplemented with appropriate amounts of antibiotics: none, 100 µg/mL carbenicillin to maintain plasmid pMMB67he, pYTK095, p1530 and derivatives thereof; 25 µg/mL chloramphenicol (CM) to maintain plasmid pMMB_IVA3_Cas_Cm or select for CM resistant MG1655; 20 µg/mL gentamicin to maintain plasmid pHERD30T and derivatives; 20ug/mL kanamycin or 15 µg/mL tetracycline to maintain pKJK5 and 50 µg/mL to select for pRSF-derivates. When appropriate, the following inducer concentrations were used: 0.2–0.3 % w/v L-arabinose and 0.1 mM isopropyl β-D-1-thiogalactopyranoside (IPTG). To propagate the virulent *E. coli* phage λ-vir, we incubated a single plaque with *E.coli* GeneHogs in 3 mL of LB containing 10 mM MgSO4 at 37 °C for 3 h. Then we added the 3 mL phage-bacteria mix to a 20 mL culture of *E.coli* GeneHogs of an optical density (OD at 600 nm, hereafter OD600) of ∼0.8, which we incubated at 37 °C for 6 h or until clear. We collected phages by sterile filtering the lysate and storing it at 4 °C over chloroform.

### p1530 transfer rate estimation

To estimate p1530 transfer rates [mL (CFU h)^−1^], we used the Approximate Extended Simonsen Model, which accounts for varying growth rates of donor, recipient, and transconjugants and estimates a time window for reliable transfer rate estimations ^69^. The p1530 plasmid transferred from its native host *K. pneumoniae* 808330 to the clinical *E. coli* Z1269 and *K. pneumoniae* SB5442, both carrying the Cm resistance-plasmid pACYC184, and to STm 14028 with the chromosomal Cm marker *marT::cat.* In brief, we grew four independent overnight cultures of donor and recipient strains supplemented with appropriate antibiotics, washed them by pelleting and resuspending, and mixed 1µL of 6.5-fold diluted donor and recipient cultures each to 150µL LB in a 96-well plate (final 1000-fold dilution). We enumerated donors, recipients, and transconjugants in mating populations after 6 h of growth without agitation and estimated their growth rates (h^−1^) based on hourly OD600 measurements (Tecan NanoQuant Infinite M200 Pro) using the R package Growthcurver. Finally, we estimated transfer rates with the R package conjugator ^69^

### Cas7 phylogenetic tree

We selected a representative set of type IV-A CRISPR-Cas systems and used CRISPRCasTyper ^65^ to extract the Cas7 (Csf2) protein sequences. The chosen type IV-A loci covered the diversity observed across the type IV-A variants (A1-A3) ^7^ and included the reference type IV-A1 systems studied in previous works ^5,10,12^. To root the subsequent tree, we used the Cas7 from a previously studied type IV-B system ^6^. Mafft (v.7.0) was used to generate the multiple sequence alignment (Gap open penalty: 1.53; Gap extension penalty: 0.123) ^70^ and an approximately--maximum-likelihood phylogenetic tree was generated with FastTree ^71^. The resulting tree was visualized with ITOL ^72^.

### Computational analysis to identify plasmids targeted by type IV-A3

Type IV-A3 CRISPR-Cas systems were retrieved by running CRISPRCasTyper ^65^ on the ENA bacterial genome database ^73^. We extracted spacers from the identified type IV-A3 CRISPR arrays and dereplicated them using cd-hit-est *(v.4.8.1) ^74^* (90% identity and 90% coverage), yielding a dataset of 450 non-redundant spacers. (**Suppl. Table S5**). To identify plasmids and phages targeted by type IV-A3 systems (i.e, carrying matching protospacers), we used the BLAST suite of programs, v.2.6.0+ ^75^ to screen the spacer queries in >55,000 plasmid and bacteriophage sequences. As a plasmid database, we employed the 21,520 plasmids retrieved from the complete genomes available in the NCBI non-redundant RefSeq database in March 2021. As a bacteriophage database, we used the 34,718 bacteriophage sequences available in the NCBI Virus Collection in June 2023. We indexed both databases with makeblastdb (default parameters) and used blastn (v.2.6.0+) to screen for matching protospacers, with the option -task blastn-short and an E-value threshold of 0.05 given the short length of the queries. Hits against type IV-A3-encoding plasmids – identified by running CRISPRCasTyper ^65^ on the PLSDB plasmid dataset ^76^ (**Suppl. Table S6**) – were discarded to avoid potential matches against type IV-A3 CRISPR arrays. Moreover, we only retrieved hits showing >95 % identity and >95 % coverage for further analysis. This resulted in a total number of 3,046 hits, 3,035 against the plasmid database, and 11 against the phage database. 58 % showed 0 mismatches in the alignment, and 42% showed one mismatch. No alignment with >1 mismatch was retrieved.

### Characterization of type IV-A3 carrying and targeted plasmids

For both, type IV-A3 carrying and targeted plasmids, we identified plasmid incompatibility groups with PlasmidFinder, v.2.0.1 ^66^, using the database of *Enterobacteriales* (v.2023-01-18), and antimicrobial resistance genes with the software AMRFinderPlus, v.3.11.4 ^77^. Additionally, some of these plasmids have previously been characterised as phage-plasmids ^78^. Finally, to characterise the mobility of plasmids we identified Mating Pair Formation (MPF) system and relaxase (MOB) with CONJScan, v.2.0.1 ^79^, and oriT with an in-house protocol previously described ^80^. The MPF, MOB, and oriT allowed us to classify plasmids as conjugative (putatively complete MPF system with a relaxase), decay conjugative (incomplete MPF system with a relaxase), MOB-mobilizable (relaxase in the absence of an MPF system), and oriT-mobilizable (presence of an *oriT* and absence of both MPF and MOB). The remaining replicons were considered as non-transmissible. To visualize this data we used the R package ggplot2, v.3.3.5 ^81^, with the addition of the R packages UpSetR, v.1.4.0 ^82^ and ggridges, v.0.5.3 ^83^ where required.

### Design and cloning of expression vectors

For the construction of expression vectors we performed USER cloning (NEB) or Gibson Assembly (NEB) following the manufacturer’s instructions. For exchanging the spacer sequences in pHerd_IV-A3_mini-array_NT we used Golden Gate DNA Assembly or restriction cloning, digesting 500 ng of the backbone with *Bsa*I-HF (NEB) and ligating 5 µL of 5 µM spacers (annealed oligos) with 80ng of *Bsa*I-digested backbone using the T4 ligase (NEB). We chose spacer sequences based on protospacer position and their association with a 5’-AAG-3’ motif and annealed them from two oligonucleotides with the according restriction site overhangs (95 °C for 5 min, 23 °C for 15 min). All constructs were then transformed into *E.coli* and constructs were confirmed by Sanger and/ or Oxford Nanopore sequencing. All oligonucleotides used in this study are listed in **Suppl. Table S7**.

### RNA Sequencing and data analysis

To test type IV-A3 activity, we analysed crRNA processing in both, *K. pneumoniae* 808330 under natural expression from p1530 and in *E. coli* MG1655 from pMMB_IVA3_Cas_CRISPR. For both we extracted small RNAs with the mirVana isolation kit (Ambion), treated with DNase I (New England Biolabs, NEB), end-repaired with T4 Polynucleotide Kinase (NEB), and submitted final products to library preparation (NEBNext Ultra RNA Library Prep Kit for Illumina), following the manufacturer’s instructions. For the transcriptomic analysis of *K. pneumoniae* 808330, we extracted total RNA with the mirVana isolation kit (Ambion), treated with DNase I (NEB), rRNA depleted with a NEBNext rRNA Depletion Kit (Bacteria), and submitted final products to library preparation using a NEBNext Ultra II Directional RNA Library Prep Kit for Illumina following the manufacturer’s instructions. We sequenced with an Illumina MiniSeq System in single-end mode, generating 150 nucleotide reads and quality control using FastQC. We trimmed reads with Cutadapt and aligned them to the genome of 808330 and *E. coli* MG1655 using Hisat2 ^84–86^. To calculate the abundance of transcripts we used the RPKM method ^87^. For data analysis, coverage plots, and scatter plots we used the R package ggplot2 ^88^. Expression and purification of the type IV-A3 ribonucleoprotein complex.

To express the type IV-A3 ribonucleoprotein complex and the crRNA, we grew overnight cultures of single colonies of *E. coli* BL21 Star containing the plasmid-encoded type IV-A3 complex with a C-terminal Gly-His6-tag at Cas7 (Csf2), each in 15 mL terrific broth (TB, Thermo) at 37 °C with required antibiotics and at 200 rpm (here and the following steps). These starter cultures we subcultured in 1 L TB with required antibiotics to an OD600 of 0.6 – 0.8, before induction of expression by adding IPTG and further growth for 3 h. We pelleted cells by centrifugation (3,600 x g, 30 min, 4 °C) and resuspended them in 20 mL lysis buffer (10 mM HEPES-Na, pH 8.0, 150 mM NaCl, 40 mM imidazole) before cell lysis by sonication using a Vibra-Cell ultrasonic processor at 40 % amplitude for 5 min with pulses of 3 s at 3 s intervals. We cleared lysates by centrifugation (47,384 x g for 20 min at 4 °C) and applied supernatants onto 1 mL HisTrap FF columns (Cytiva, pre-equilibrated in lysis buffer, for Ni-NTA affinity chromatography at 4 °C). After a wash step with 15 column volumes of lysis buffer, we eluted proteins with three column volumes of elution buffer (10 mM HEPES-Na, pH 8.0, 150 mM NaCl, 500 mM imidazole). We concentrated proteins to 0.5 mL at 4 °C and further purified by size-exclusion chromatography using a Superose 6 Increase 10/300 GL column (Cytiva, equilibrated in size exclusion buffer: 10 mM HEPES-Na, pH 7.5, 150 mM NaCl) at 4 °C. We then concentrated main peak fractions to 0.5 mL at 4 °C and estimated concentrations based on the absorbance at 280 nm using a NanoDrop Eight spectrophotometer (Thermo) and extinction coefficients based on an assumed Cas protein complex stoichiometry of 1:1:6:1 (Cas5:Cas8:Cas7:Cas6).

### Verification of *in trans* spacer acquisition

A similar approach to detect the acquisition of synthetic protospacers has been used by Shipmann et al., 2016 ^16^. In brief, we grew *E. coli* MG1655 carrying p1530 in combination with either pHerd_Cas1/2e or pHerd30T_ev (negative control) overnight, each in triplicates, and subcultured cells in LB containing 0.2 % w/v L-arabinose until they reached an OD600 of ∼0.4. We made each replicate electrocompetent according to standard laboratory procedures and electroporated 100 ng of the double-stranded protospacer PS (PSA33 ^16^) as annealed oligonucleotides. As a negative control, we electroporated cells without any PS DNA. Cells were recovered for 1.5 h in LB supplemented with 0.2 % w/v L-arabinose, spun down and resuspended in 50 µL water, and stored at 4 °C until further processing. To confirm the acquisition of the new PS in the type IV-A3 array, we performed two PCRs on each of these templates: one PCR to amplify the leader-proximal end (pFB29, annealing to PS/pFB39) and one to amplify the leader distal end (pFB28, annealing to PS/pFB74). We verified spacer integration in the leader-repeat junction by agarose gel electrophoresis and PCRs yielded amplicons of 108 bp and 328 bp, respectively. We subjected PCR products from each replicate to Sanger-sequencing twice, once with pFB29 (leader-proximal end) and once with pFB28 (leader-distal end). We performed PCRs with the Phusion DNA Polymerase (Thermo) following the manufacturer’s protocol and with an annealing temperature of 60.1°C for 35 cycles.

### Detection of spacer acquisition in native CRISPR arrays

To facilitate genome-wide spacer acquisition we used a p1530Δ*dinG* (NT) containing *K. pneumoniae* 808330. We generated this mutant in *E. coli* GeneHogs using the lambda red recombinase system ^89^. For the assay, we conjugated p1530Δ*dinG* back into *K. pneumoniae* 808330, from which we previously cured p1530. We grew the p1530Δ*dinG* containing *K. pneumoniae* 808330 from single colonies with pHerd_Cas1/2e or pHerd30T_ev (negative control) in quadruplicates in LB supplemented with L-arabinose (0.2 % w/v) overnight. For each sample, we extracted total DNA with the DNeasy Blood &Tissue Kit (QIAGEN) from 1.5 mL cultures, which we used as templates for subsequent PCR reactions (100 ng). To monitor adaptation, we amplified the leader-proximal end of the CRISPR array by CAPTURE PCR ^20^: a first PCR with primers targeting the leader (pFB88) and the first spacer (pFB89, spacer1) isolated the leader-proximal end (leader-repeat1-spacer1 for unextended and leader-new repeat-acquired spacer-repeat1-spacer1 for extended arrays). We separated extended amplicons from unexpanded products by agarose gel electrophoresis (2.5 % w/v), cut invisible bands of elongated arrays (174 bp), and isolated DNAs with the GeneJET Gel Extraction Kit (Thermo). These served as templates for a second PCR: we amplified extended arrays with pFB88 and degenerate primers targeting repeat1, whose 5’ ends are not complementary to the 3’ end of the leader (adenine, pFB90, pFB91, pFB92). Importantly, this method introduces a bias because spacers that carry the base adenine at their 3’ end are likely not amplified. After this selective PCR, we separated expanded/unexpanded amplicons by agarose gel electrophoresis (1.5 % w/v). One replicate did not show a band the size of expanded arrays and was dismissed. We performed PCRs with the Phusion DNA Polymerase (Thermo) following the manufacturer’s protocol and with an annealing temperature of 67.2 °C for 35 cycles. To reach a high enough DNA concentration for high-throughput sequencing, we performed each PCR with multiple reactions for each sample. Array-amplicons were sequenced at Novogen (Illumina NovaSeq, 150 nucleotides paired-end reads, UK) after adaptor ligation (NEBNext Ultra II DNA Library Prep Kit for Illumina). After sequencing, we de-multiplexed samples by index and had an average of 4,616,517 read pairs. First, we filtered and trimmed reads according to base qualities with Trimmomatic (v0.39) ^90^ with the following parameters: PE LEADING:5 TRAILING:5 MINLEN:80 AVGQUAL:20. Filtering removed on average 0.11% of the read pairs. Second, we merged forward and reverse reads with PEAR (v0.9.6) ^91^ with default parameters. On average 98.35 % of filtered read pairs were merged. To extract spacers we used cutadapt (v1.18) ^84^ with the pattern (partial leader + *repeat* + spacer(…) + *partial repeat*, or the reverse complement) with the -g option: "GCTGGTGGATTTTAGTGGCGCTATTTAATATTTTATAATCA-ACCGGTTATTTTTAGA*GTATTCCCCCCGTGTGCGGGGGTTATCG*…*GTATTCCCCCCGTGTGCG*". From on average 93.4 % of merged read pairs a spacer of any length could be extracted, of which 97 % were between 31 and 35 bp. We reverse-complemented spacers extracted with the reverse complement pattern to match the transcribed strand. To align spacers we searched for perfect matches on either strand of present genomes and excluded matches to the array on p1530 and spacers aligning multiple locations; this resulted in matches for 78.5 % of the 33 bp spacers, which we used for the SAM analysis.

### pKJK5 targeting assays

To test the type IV-A3’s capacity to target the environmental and tetracycline-resistant IncP-1 plasmid pKJK5 tagged with *gfp* ^24^ in *E. coli*, we overexpressed the IV-A3 Cas operon from the P_TAC_ promoter and complemented it with a crRNA (mini array; repeat-spacer-repeat sequence) expressed from the P_Bad_ promoter crRNAs contained spacers targeting *trfA*, *parA*, *traI*, and *gfp* in the coding (+) and template strand (-), and the *oriT*, and we used the non-targeting pHerd_IV-A3_mini-array_NT as a negative control. Plasmids are given in **Suppl. Table S4** and explanations for a spacer design are provided in **Suppl. Fig. S7A-C**.

With this, we set up three independent targeting assays, two to test type IV-A3 interference with pKJK5 transfer and one to test IV-A3 interference with pKJK5 stability within a bacterial population. In the pKJK5 incoming (from D0 to RIV-A3) (**Fig. 3B**) and the pKJK5 outgoing (from DIV-A3 to R_0_) (**Fig. 3D**) assays, we estimated IV-A3 interference in the *E. coli* MG1655 recipient R_IV-A3_ and donor D_IV-A3_, respectively, as the pKJK5 transfer efficiency (CFU_transconjugants_/ (CFU_recipients_+CFU_transconjugants_), (T/(R+T)) relative to the pKJK5 transfer efficiency in the NT control. Note that for the outgoing assay, the recipients also encoded a type IV-A3 and crRNA to avoid obscuring the interference signal by secondary pKJK5 transfer (from transconjugants to recipients) and a chromosomal Cm marker. In brief, after overnight growth of recipient and donor strains in biological triplicates and with required antibiotics, we diluted 1:100 and re-grew strains in LB + inducers and appropriate antibiotics into exponential phase (∼3 h). We removed antibiotics by spinning and resuspending cultures in LB + inducers and incubated at RT for 15 min. To initiate mating cultures we spun 500 µL each, resuspended in 20 µL LB + inducers, and mixed 20 µL of donor and recipient to 40 µL mating cultures which were allowed to conjugate on LB agar plates + inducers for 3h after drying. We rescued mating drops with a loop and resuspended them in 500 µL LB prior to dilution plating on LB agar plates + inducers and appropriate antibiotics to select either donors+transconjugants, recipients+transconjugants, or transconjugants only. The outgoing assay differed only in that conjugation time was limited to 35 min and inducers were added while growing cultures to the exponential phase. It was not possible to evaluate *trfA* targeting, as pKJK5 cannot be stably maintained in the donor strain under type IV-A3 induction.

For the plasmid stability assay, we grew *E. coli* MG1655 encoding the type IV-A3 Cas operon and crRNAs in four biological replicates overnight in a randomized 96-well plate containing 150 µL LB with required antibiotics and 0.2 % w/v glucose to inhibit crRNA expression. To initiate the stability assay (day 0) we twice spun cultures to remove glucose and resuspended in 150 µL LB + inducers and required antibiotics. We grew cultures for 4 days with daily passaging of 1.5 µL of grown cultures into fresh medium and plating on LB agar plates + inducers and appropriate antibiotics to enumerate plasmid-carrying and plasmid-free subpopulations. To perform statistical analyses we used the detection limit of 100 CFU/ mL when we did not obtain any plasmid-carrying colonies under *trfA* targeting.

To measure population growth rates under type IV-A3 targeting and for the NT control we grew five biological replicates of the strains used above as donor D_IV-A3_ in LB + required antibiotics for vector and pKJK5 selection overnight. To initiate the experiment we diluted cultures 100-fold by adding 1.5 µL to a randomized 96-well plate containing 150 µL of fresh LB + antibiotics and inducers. Cultures grew for 20 h in a Tecan NanoQuant Infinite M200 Pro and were shaken prior to the hourly measurement.

### *gfp* targeting in transconjugants

After the incoming pKJK5 plasmid targeting assay, we measured green fluorescence signal of transconjugants. For each replicate of the strains targeting *gfp* in the coding (+) and template (-) strand, and the NT control, we diluted the cells in 3 mL PBS and proceeded to analyze them through flow cytometry (FACSAria Illu Becton Dickson Biosciences, San Jose, CA, USA) using a 70 μm nozzle and sheath fluid pressure of 70 lb/in2. GFP was excited by a 488 nm laser (20 mW) and detected on the fluoresceine isothiocyanate A (FITC-A) channel; bandpass filter of 530/30 nm. mCherry was excited with a 561 nm laser (50 mW) and detected on the phosphatidylethanolamine (PE)-Texas Red-A channel; bandpass filter of 610/20 nm. We set detection thresholds to 200 for forward (FSC) and side scatter (SSC) and used the BD FACSDiva software (v6.1.3) for data analyses. Briefly, we used scatterplots of particle FSC vs. SSC to delimit gates for bacterial events, thus excluding background noise. We used bivariate contour plots (FITC vs. PE-Texas Red) to gate GFP- and mCherry-positive bacterial cells. We used cell counts of 1000 to 3000 threshold events/ s, processed at flow rate 1, and recorded a total of 30,000 bacterial events for each replicate. We enumerated the following fluorescent phenotypes, i) total cells expressing mCherry (red), ii) total cells expressing *gfp* from pKJK5 and mCherry (red-and-green). We then calculated the percentage of cells that were red-and-green (transconjugants) of the total red population (recipients).

### Phage targeting assay

We assessed the functionality of type IV-A3 in phage targeting through phage-spotting assays, by evaluating the replication of CRISPR-targeted phage λ-vir on bacterial lawns (GeneHogs) in comparison to the NT control. In brief, we overexpressed the IV-A3 Cas operon under *Ptac* promoter supplemented with crRNAs (targeting and NT) expressed from *PBad* promoter overnight in *E. coli* GeneHogs in triplicates. We mixed 150 µL of bacterial overnight cultures with 4 mL of molten top agar (0.7 % w/v) supplemented with 10 mM MgSO4, L-arabinose (0.3 % w/v), and IPTG. We poured the mix onto LB agar plates containing MgSO4, L-arabinose, and IPTG and spotted 4 or 10 µL of 10-fold serial diluted phage lysates onto the lawn. Plates incubated at 30 °C to count PFU the next day.

### Isolation of escaper phages

To isolate phages that escaped CRISPR-targeting, we spotted 20 ul of undiluted ancestor phage on the lawn of the respective targeting strain and stroke out the phage across the plate, and incubated them overnight at 30 °C. We then picked single plaques of spontaneous escapers and restreaked them on a new lawn. We repeated this single plaque isolation three times to ensure that no mixed genotypes of phages remained. We then amplified the targeted sites via PCR and Sanger sequenced the fragments for escapers in gene B and the intergenic region. We could not get a PCR product for the R escaper and therefore extracted the total DNA from the escaper and the ancestor λ-vir phage lysate after enrichment for high phage titer (> 10^7^ pfu/ mL) DNA with the DNeasy blood and tissue kit (QIAGEN), starting from the Proteinase-K treatment step to lyse the phages ^92^. We prepared sequencing libraries using the Illumina NEXTERA XT Kit for tagmentation and amplification (12 cycles), following the manufacturer’s instructions and purified libraries using the bead-based HighPrep™ clean-up Kit (MagBio Genomics). Paired-end sequencing was performed on an Illumina MiSeq platform using Miseq V3 chemistry (2 x 300 cycles), according to the manufacturer’s protocol.We used CLC Genomics Workbench (v20.0.4) for adapter trimming and generation of de novo assemblies and annotated assembled genomes with Rapid Annotations (Subsystems Technology tool kit (RASTtk) accessed through PATRIC (v3.6.12) ^93^. To determine the molecular mechanisms of escaping, we aligned escaper assemblies to the ancestral reference with snapgene (v6.0.2). With NCBI nucleotide blast (standard parameters), we identified genomic region acquired by escapers to be also located in the chromosomes of laboratory *E.coli* as (DH10B, DH5alph) from which also GeneHogs descend (e.g., NCBI: CP000948.1). We visualized phage-λ genomes with clinker using standard parameters ^94^.

### TXTL-based PAM Assay: PAM-DETECT

To identify the preferred TIM sequence for type IV-A3 target interference, we used the PAM-DETECT methodology, as described previously ^30^. Briefly, we used a vector-based protospacer library with randomised TIMs. Protospacers containing a functional TIM sequence are protected from cleavage, since successful IV-A3 complex binding to the protospacer prevents plasmid cleavage in a subsequent restriction digestion step. Enrichment of TIMs is then assessed via high-throughput sequencing. We used a plasmid containing a library of five randomized nucleotides (potential TIMs) flanked by a protospacer containing a PacI restriction site and expressed the type IV-A3 Cas components and crRNA from separate plasmids. The 6 µl TXTL reaction consisted of 3 nM plasmid encoding for the IV-A3 effector complex, 1 nM crRNA-encoding and PAM-library plasmid each, 0.2 nM T7 RNA polymerase, 0.5 mM IPTG, and 4.5 μL myTXTL Sigma 70 Master Mix. For the negative control we replaced the IV-A3 encoding plasmid with an equal volume of water. We incubated TXTL reactions at 29 °C for 6 h and digested at 37 °C with PacI (NEB R0547S) as instructed by the provider and added water instead of PacI for the undigested control. After PacI inactivation, we added 0.05 mg/ mL Proteinase K (Cytiva), incubated at 45 °C for 1 h and after Proteinase K inactivation extracted the remaining plasmids with standard EtOH precipitation.

For NGS library preparation we added adapters and unique dual indices in a two-step amplification process using the KAPA HiFi HotStart Library Amplification Kit (KAPA Biosystems, KK2611) and purified samples with Agencourt AMPure XP (Beckman Coulter, A63881). We used a Illumina NovaSeq 6000 (paired end, 2 × 50 bp, 2 million reads per sample) sequencer and for NGS data analysis we followed ^95^ and ^96^. First, we normalized the read counts of every TIM with the total number of reads and calculated the ratio of digested to undigested sample reads. The sum of ratios for a given nucleotide at a given position we then divided by the sum of the ratios of all nucleotides at that given position (resulting in 25 % in case of no enrichment/depletion). Finally, to assess the amount of library plasmid protected from restriction digestion, we performed qPCR using the SsoAdvanced Universal SYBR Green Supermix (Biorad, cat#1725271) with primers amplifying a 100 bp spanning the *Pac*I recognition site of the library plasmid and primers amplifying a 100 bp region on the T7 RNA polymerase encoding plasmid as a control. We quantified reactions with the QuantStudio Real-Time PCR System (Thermo Fisher Scientific) and an annealing temperature of 68 °C, according to manufacturers’ instructions.

### *mCherry* chromosome targeting

To shed light on the relevance of DinG for target interference through transcriptional repression, we used three type IV-A3 variants (wildtype, DinG knockout mutant Δ*dinG*, catalytically inactive DinG mutant *dinGmut*) in an assay targeting the chromosomally encoded *mCherry* or its promoter *P_lpp_*. We grew four biological replicates of MG1655 for each possible combination of the IV-A3 variant and crRNA (three variants x eight crRNAs and the NT control, 27 in total) overnight with appropriate antibiotics in a 96-well plate. To initiate the experiment, we pin-replicated ∼1 µL of each culture into a black (transparent bottom) 96-well plate containing 150 µL LB + appropriate antibiotics + inducers. Cultures grew for 24 hours in a Tecan NanoQuant Infinite M200 Pro. We measured OD600 and fluorescence intensity (Excitation 582 nm, Emission 620 nm) in an interval of 15 min. For each replicate and time point, we normalized the mCherry signal with the corresponding OD600 measure and selected timepoint 15 h for further analysis. To identify the best fit model explaining most of the observed variation in mCherry signal, we performed linear regression models with the three predictors IV-A3 variant, position (promoter vs. gene), and strand with interactions.

### Bio-layer interferometry of the IV-A3 complex on double-stranded DNA

To test type IV-A3 CRISPR-Cas complex affinity to double-stranded DNA targets, we performed bio-layer interferometry experiments in size exclusion buffer corresponding to the respective analyte on an Octet K2 System (Pall ForteBio). We prepared the target dsDNA ligand by annealing oligonucleotides BTS and NTS at a ratio of 1:1.5 (5 µM BTS: 7.5 µM NTS) in size exclusion buffer by heating the annealing reaction at 95 °C for 5 min and slowly cooling it down to RT. To immobilize the dsDNA ligand on an Octet SAX 2.0 biosensors (Sartorius) we prepared 200 μL of a 25 nM dsDNA solution in a black 96-well plate and performed a loading step for 120 s followed by a washing step for 30 s. We diluted the IV-A3 complex, either with NT or targeting crRNA, from 1 µM to 25 nM in a 200 μl final volume dilution series and measured association and dissociation for 300 s and 180 s, respectively. The baseline was recorded prior and after association/dissociation for 30 s in the matching size exclusion buffer and for each measurement. A reference omitting the dsDNA from the solution was recorded to subtract reference curves from sample curves. For K_D_ determination the reference-subtracted binding and dissociation curves were fit to the standard 1:1 local binding model using the Pall ForteBio analysis software.

### Cell-free transcription-translation assays

To further explore the targeting activity of the distinct type IV-A3 variants observed in our *in vivo* transcriptional repression analysis, we performed a cell-free transcription-translation (TXTL) assay. We used individual plasmids encoding one of the three type IV-A3 variants (wildtype, Δ*dinG*, *dinGmut*), deGFP, and one of the *degfp*-targeting crRNAs (one on each strand of PT7, four on each strand within the *degfp*; 10 in total and a NT control). Because the type IV-A3 *cas* and *degfp* are under the control of a PT7, we also included a plasmid encoding the T7 RNA polymerase. To assess the targeting activity of the IV-A3 variants at distinct interference loci, we assembled the following reaction mixes using an Echo525 Liquid Handling system (Beckman Coulter, 001-10080): We added each possible combination of the IV-A3 variant and crRNA and the remaining plasmids to the myTXTL mix (Arbor Biosciences, 507025-ARB) at the following final concentrations: 2 nM each for plasmids encoding type IV-A3 and the crRNA, 1 nM of the deGFP plasmid and 0.2 nM of the T7 RNA polymerase plasmid (3 µl per reaction mix and four replicates each). To measure background fluorescence, we assembled an additional mix that only contained myTXTL mix and water. We incubated these reactions at 29 °C for 16 h in a plate reader (BioTek Synergy Neo2) and measured fluorescence every 3 minutes. We plotted endpoint measurements as the RFU after subtracting the background fluorescence from each reaction.

### Re-sensitizing of antibiotic-resistant strains

To investigate type IV-A3’s capacity to re-sensitize bacterial strains by transcriptional repression of β-lactamases we first re-sensitized *E. coli* DH10B by targeting *bla*_CTX-M15_ encoded on the clinical *E. coli* plasmid p1ESBL. In brief, we grew four biological replicates of targeting and NT strains, differing only in crRNA, overnight with appropriate antibiotics, and the next day we diluted 1:100 and re-grew strains in 2 mL LB + inducers and appropriate antibiotics into exponential phase (∼3 h). We spun and resuspended cultures in 80 µL LB, serially diluted, and plated them on LB agar + inducers and appropriate antibiotics to select either p1ESBL-carrying cells or *bla*_CTX-M15_ expressing cells. To be able to enumerate plasmid-carrying cells independent of *bla*_CTX-M15_ expression we inserted a Kan marker on p1ESBL.

Second, we re-sensitized the clinical *K. pneumoniae* 808330 (Δp1530) by targeting the chromosomal *bla*_SHV-187_ with a combination of broth microdilution and targeting assay. We followed the 96-well plate-based MIC assay protocol from Wiegand et al. ^97^ but made several modifications to accommodate IVA3-interference during the assay, such as adding appropriate antibiotics for type IV-A3 and crRNA selection and inducers. Further, we substituted the Mueller-Hinton growth medium with LB. We used the same Cas operon- and mini-array setup as described for the plasmid targeting above but replaced the Amp cassette on the IV-A3 encoding pMMB67he with a Cm marker. We grew four biological replicates for strain 808330 carrying type IV-A3 and either a crRNA or the NT control overnight with appropriate antibiotics. To initiate the MIC/ targeting assay, we diluted 1:100 and re-grew strains in LB + inducers and appropriate antibiotics into exponential phase (∼3 h). We measured OD600 of cultures, adjusted them to an OD600 = 0.065, and diluted 1:100 to reach a final OD600 = 0.00065 in LB + 2-fold required concentrations of inducers and the antibiotics Cm and Gm for vector selection. To assay the MICs we added 50 µL per culture to the 96-well plate containing 50µL LB per well + specific Amp concentrations (10 concentrations with two-fold reduction steps from 32-0.0625 mg/L). Further, to compare the type IV-A3 mediated MIC reduction to the one achieved by a *bla*_SHV-187_ null-mutant we performed a MIC assay as above with 808330 (Δp1530) and the same strain with an additional Δ*SVH* mutation. Here no subculturing step prior to the MIC assay and no inducers or antibiotics additional to Amp were required. We interpreted MIC breakpoints for Amp according to EUCAST guidelines (v4.0, 2022-01-01) after 18-24 h of static growth ^98^. To compare the IV-A3 mediated MIC reduction to the one achieved by the addition of the β-lactamase inhibitor clavulanic acid we performed a disc diffusion assay under targeting using Amp (10 µg) and amoxicillin-clavulanic acid (Amc; 20/10 µg). We grew four biological replicates for strain 808330 (Δp1530) carrying type IV-A3 and either a crRNA or the NT control overnight + inducers and appropriate antibiotics. We adjusted strains to OD600 = 0.02 and distributed 2 mL thereof on LB agar plates, removed excess liquid by pipetting, and placed the filters on plates after drying. After 20 h incubation, we measured the diameters of inhibition zones by hand and interpreted MIC breakpoints according to EUCAST guidelines (v13.0, 2023-01-01; ^99^.

## ACKNOWLEDGMENT

We thank the Synthetic Biology and Microbial Evolutionary Genomics groups at Institut Pasteur, the Section of Microbiology at the University of Copenhagen, the Phi Lab (Peter Fineran’s group) at Otago University, and the Pathogen Ecology group at ETH Zürich for helpful discussions and mimosas. We thank David Mayo-Muñoz for valuable feedback on the manuscript. We thank Sylvain Brisse (Institut Pasteur) for providing *K. pneumoniae* strain SB and his group, especially Carla Rodrigues and Chiara Crestani, for help with Oxford Nanopore sequencing. Plasmids encoding P70A-deGFP and P70A-T7RNAP were kindly provided by Prof. Vincent Noireaux. We thank the Šikšnys laboratory (Vilnius University) for providing access to the Octet K2.

## STATISTICAL ANALYSES AND FIGURE PREPARATION

We performed all statistical analyses using R (v4.1.0) and figures were plotted with Prism9 and edited with Adobe Illustrator.

## AUTHOR CONTRIBUTIONS

R.P.-R., F.B, and S.C.-W. conceived the project.

J.V.G.F. performed RNA-seq, K.W. and F.E. performed TXTL and TIM assays, A. C. identified *K. pneumoniae* 808330. R. Č. expressed and purified ribonucleoprotein complexes and performed BLI. The other experiments were performed by F.B., S.C.-W, J.K., and S. G..

F.B., S.C.-W., K.W., F.E., J.V.G.-F., J.K., S. G., and L.M. verified the overall reproducibility of results and other research outputs.

M. A.-A. and J.R. implemented bioinformatic analyses.

M. A.-A., J.R., F.B., M.A.S, R.P.-R., and S.C.-W. performed computational analyses.

J.R., F.B., and S.C.-W. analyzed and synthesized study data by applying statistical, mathematical, and computational techniques.

R.P.-R, F.B., and S.C.-W. curated data.

R.P.-R., F.B., and S.C.-W. wrote the initial draft of the manuscript and designed figures.

All authors revised and crucially contributed to the current draft of the manuscript.

R.P.-R. managed and coordinated research activity planning and execution.

R.P.-R., P.P., C.B., D.B., A.H., S.J.S., J.S.M., L.R., and E.R. provided mentorship and oversight of the research activities.

Funding acquisition by R.P.-R, F.B., A.H., J.S.M., D.B., C.B., S.J.S, P.P., and L.R.

## SUPPLEMENTARY INFORMATION

### Supplementary Tables

**Table S1:** Spacer-protospacer analysis *K. pneumoniae* 808330 Type IV-A3 and I-E.

**Table S2:** Plasmids in *K. pneumoniae* 808330.

**Table S3.** Strains and phages used in this study.

**Table S4:** Plasmid constructs used in this study.

**Table S5:** Type IV-A3 dereplicated spacers.

**Table S6:** Type IV-A3 carrying plasmids.

**Table S7:** Oligonucleotides and DNA fragments used in this study.

The above tables are in a separate datafile and can be made available upon request.

### Supplementary Figures

**Figure S1:**
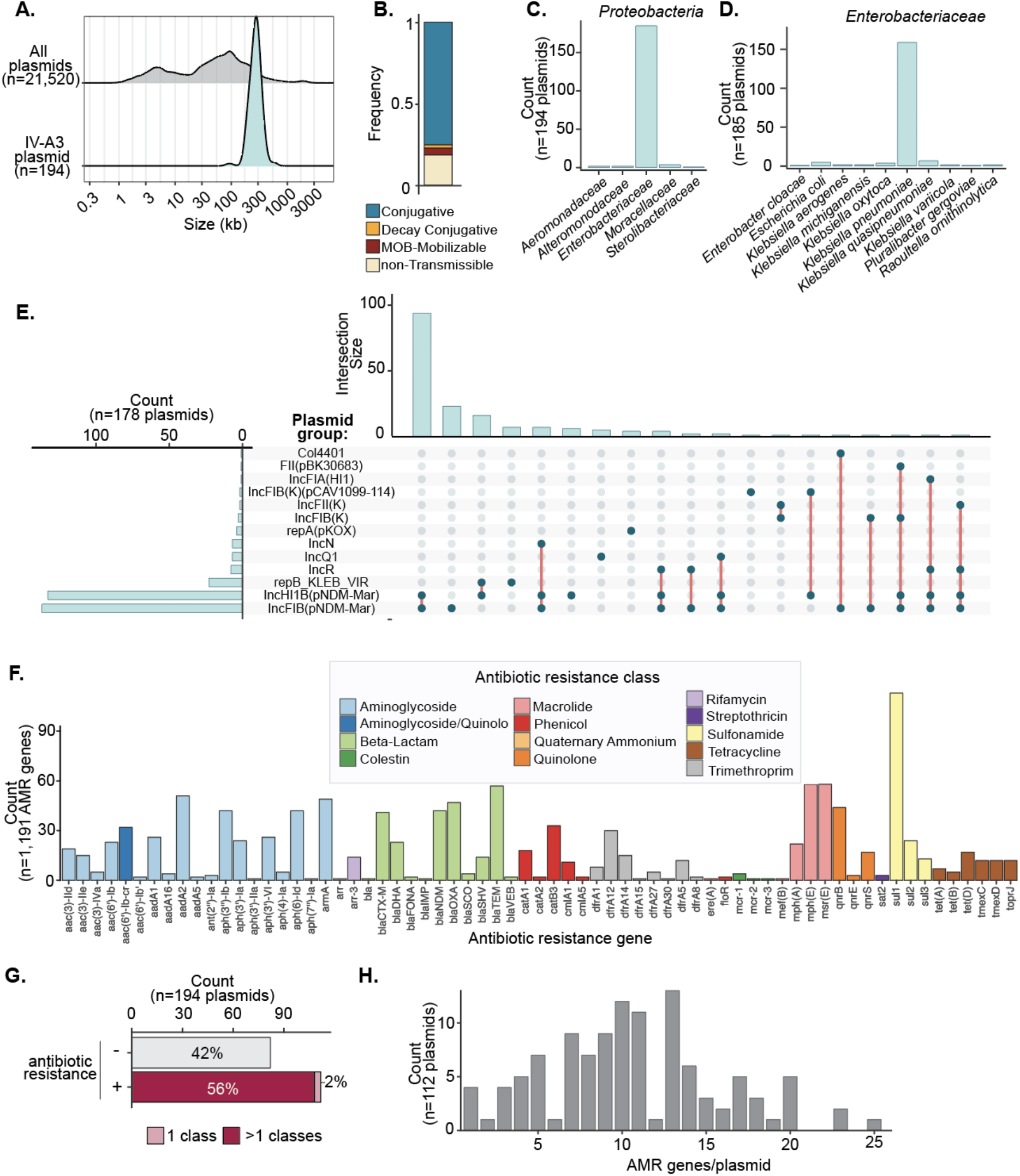
Characteristics of plasmids carrying type IV-A3 CRISPR-Cas. **A)** Size distribution of all RefSeq plasmids (top) and type IV-A3 carrying plasmids (bottom) using a Kernel density estimation. **B)** Mobility prediction of plasmids carrying type IV-A3. **C)** Family distribution of *Proteobacteria* hosts carrying type IV-A3 CRISPR-Cas systems. **D)** Species distribution of *Enterobacteriaceae* hosts carrying type IV-A3. **E)** Type IV-A3 carrying plasmids broken down by predicted plasmid replicon group. The upset plot shows the counts per group (left), and the intersection size of replicon combinations (right). Out of the 194 plasmids, 16 are not shown in this analysis (non-typeable). **F)** Distribution of antibiotic-resistance genes encoded by type IV-A3 carrying plasmids, colored by corresponding antibiotic-resistance class. **G)** Antibiotic resistance carriage by plasmids encoding type IV-A3. Counts of plasmids carrying more than five resistance genes are shown. The relative frequencies of each category are indicated inside the bars. **H)** Histogram showing the number of antibiotic resistance genes carried by type IV-A3 encoding plasmids. Only plasmids encoding one or more antibiotic-resistance genes are shown.

**Figure S2:**
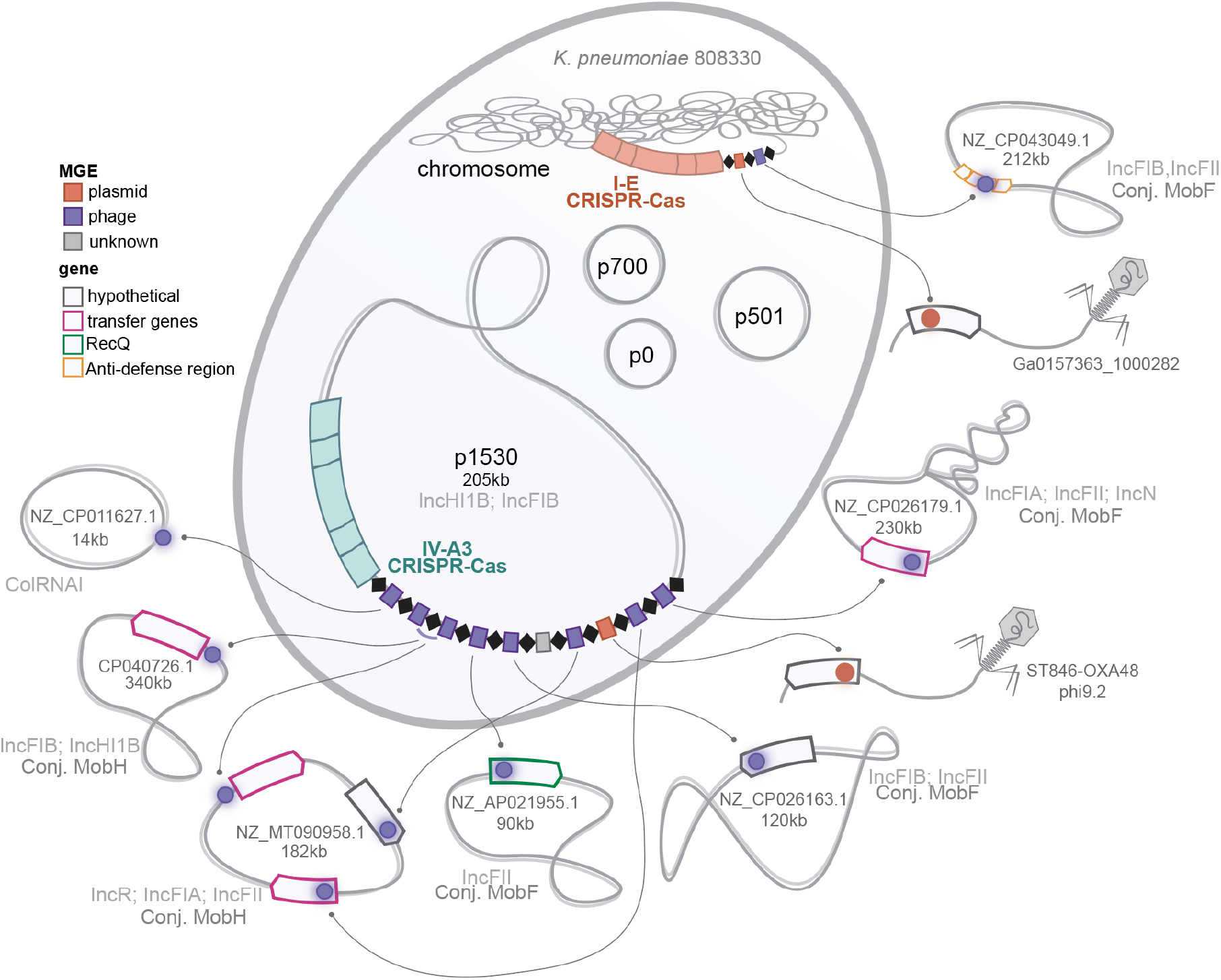
Schematic of the *K. pneumoniae* 808330 model strain highlighting the predicted targets of its CRISPR-Cas systems. Spacer-matching plasmid sequences are depicted in purple, while those targeting phages are represented in orange. The predicted function of the targeted phage- and plasmid-derived genes are indicated by colored frames. The information provided includes the names and accessions of example targeted plasmids and phages, along with the predicted Inc/Mob plasmid classifications. Additional plasmid details are summarized in **Suppl. Table S1**.

**Figure S3:**
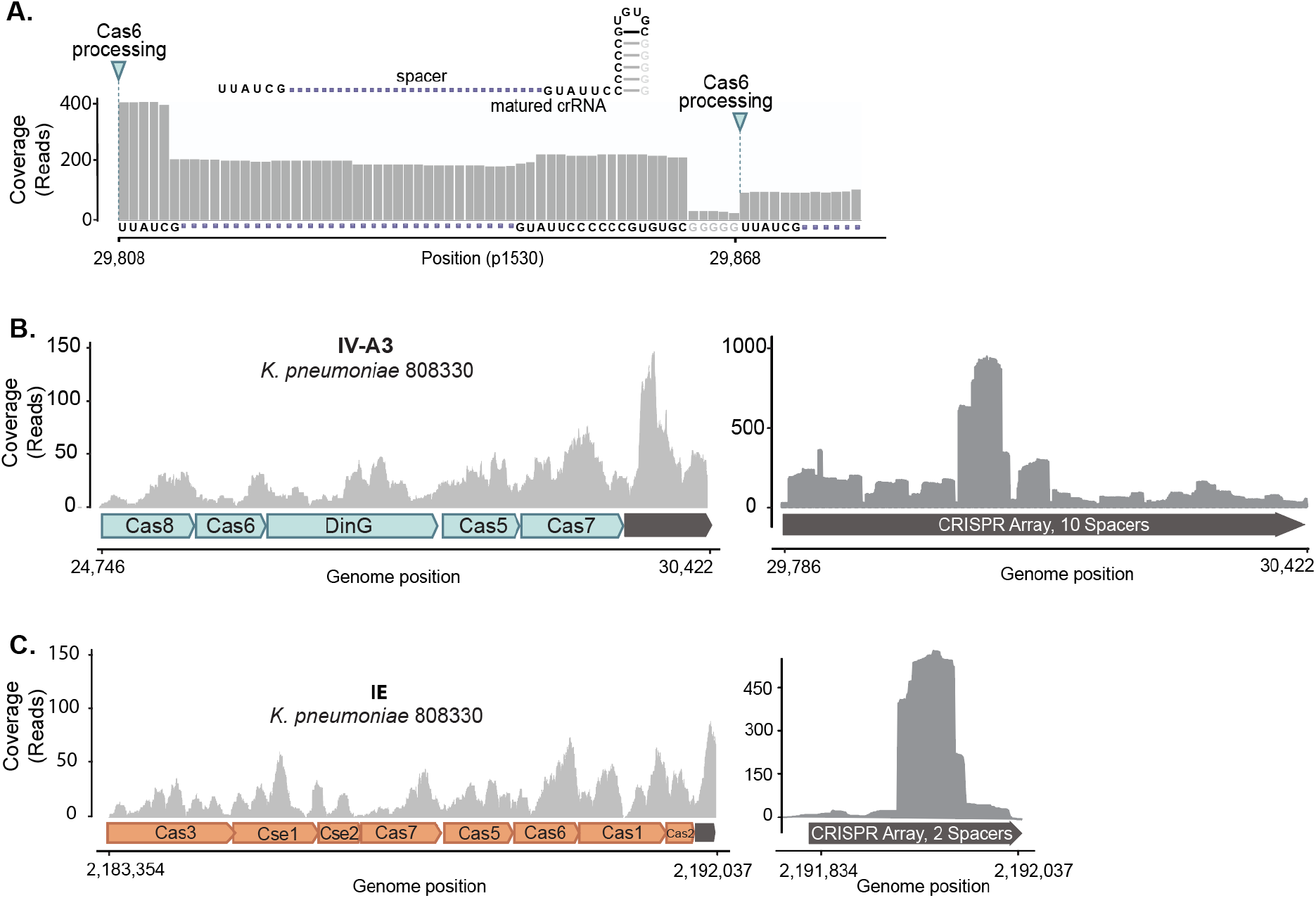
Expression analysis of the CRISPR-Cas systems encoded in *K. pneumoniae* 808330. **A)** Zoom in on small RNA-seq of mature crRNAs in *E. coli* reveals processing at the base of the extended hairpin and resulted in matured crRNAs with a six nucleotide 5ʹ handle and a 3ʹ handle comprised of 17 nucleotides, with the last four guanines of the stem missing. Schematic of the matured crRNA secondary structure is indicated as well as the Cas6-processing sites. **B)** RNA-Seq analysis of the IV-A3 CRISPR-Cas expression in *K. pneumoniae* 808330. Total RNA-seq of the type IV-A3 CRISPR-Cas locus on the left, small RNA-seq of the processed crRNAs mapped back to the CRISPR array on the right. **C)** RNA-seq analysis of the I-E CRISPR-Cas expression in *K. pneumoniae* 808330. Total RNA-seq of the type I-E CRISPR-Cas locus on the left, small RNA-seq of the processed crRNAs mapped back to the CRISPR array on the right. Coverage plots in **B** and **C** show the average of three biological replicates.

**Figure S4:**
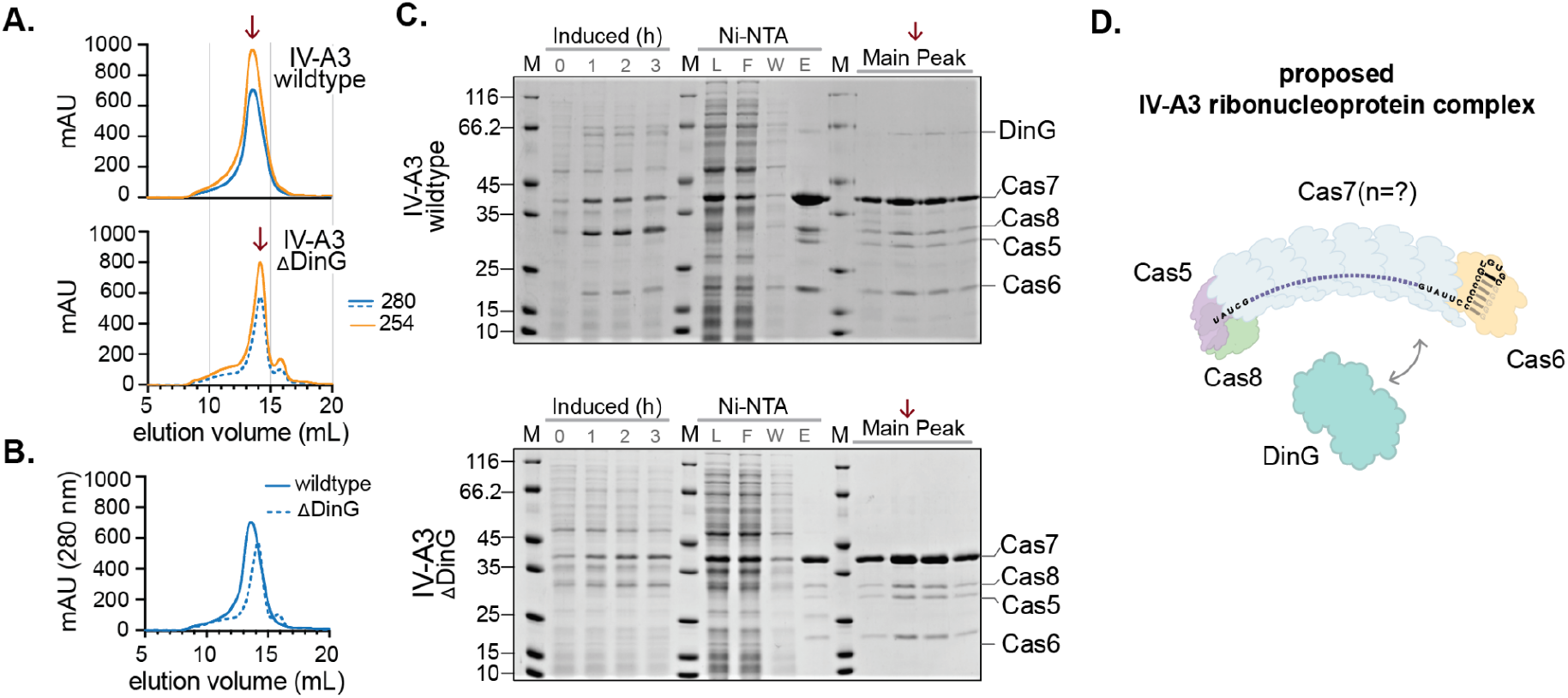
Heterologous expression and purification of the type IV-A3 ribonucleoprotein complex. **A)** Size exclusion chromatography (SEC) traces recorded at 254 nm (yellow) and 280 nm (blue) for the wildtype complex (top) and the DinG knockout version (ΔDinG; bottom), both carrying a C-terminal Gly-His6-tag at Cas7. Arrows indicate the main fraction used in the SDS-PAGE in **C**. **B)** SEC traces at 280 nm. Same data as in **A** but overlayed. **C)** SDS-PAGE for the IV-A3 wildtype (top), and IV-A3 ΔDinG (bottom) complex expression and purification, showing the whole cell content after induction (induced) at t (h) = 0, 1, 2, and 3; the Ni-NTA His-tag affinity chromatography fractions Load (L), Flowthrough (F), Wash (W) and Elution (E); and SEC main peak fractions (arrow). **D)** Schematic of type IV-A3 ribonucleoprotein complex based on in **C** detected Cas components.

**Figure S5:**
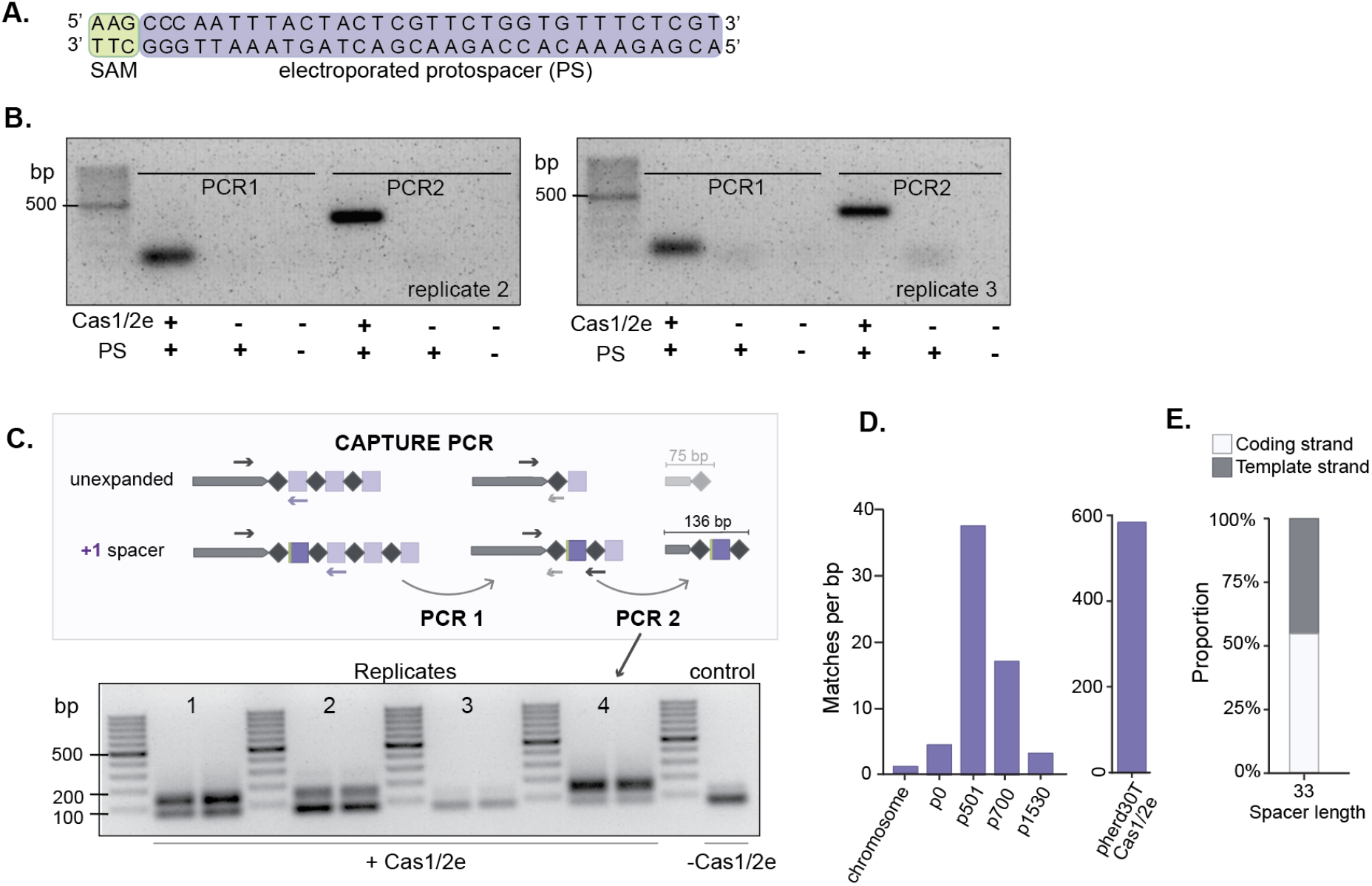
Further characterization of spacer acquisition in type IV-A3 CRISPR loci. **A)** Depiction of the protospacer used to electroporate *E. coli* MG1655 carrying the p1530-encoded type IV-A3 CRISPR-Cas system facilitating observation of Cas1/2e dependent adaptation. **B)** Replicate 2 and 3 of the amplicons confirming the sequence-specific acquisition of electroporated protospacers in the leader-repeat junction from Figure 2B. **C)** Top: Schematic showing the workflow of CAPTURE PCR comprised of PCR 1 and PCR 2. Bottom: Replicates 1-4 of amplicons of PCR 2 used for detection of Cas1/2e dependent genome-wide spacer acquisition in *K. pneumoniae*. Amplicons of 136 bp indicate spacer acquisition and amplicons of 75 bp result from secondary primer binding. Replicate 3 did not show a 136 bp amplicon and was therefore excluded from further analysis. **D)** Spacers identified from genome-wide acquisition assay mapped back to replicons present in *K. pneumoniae* 808330. The values are normalized to the size of the respective replicon. **E.** Template and coding strand acquisition for spacers of length 33 bp.

**Figure S6:**
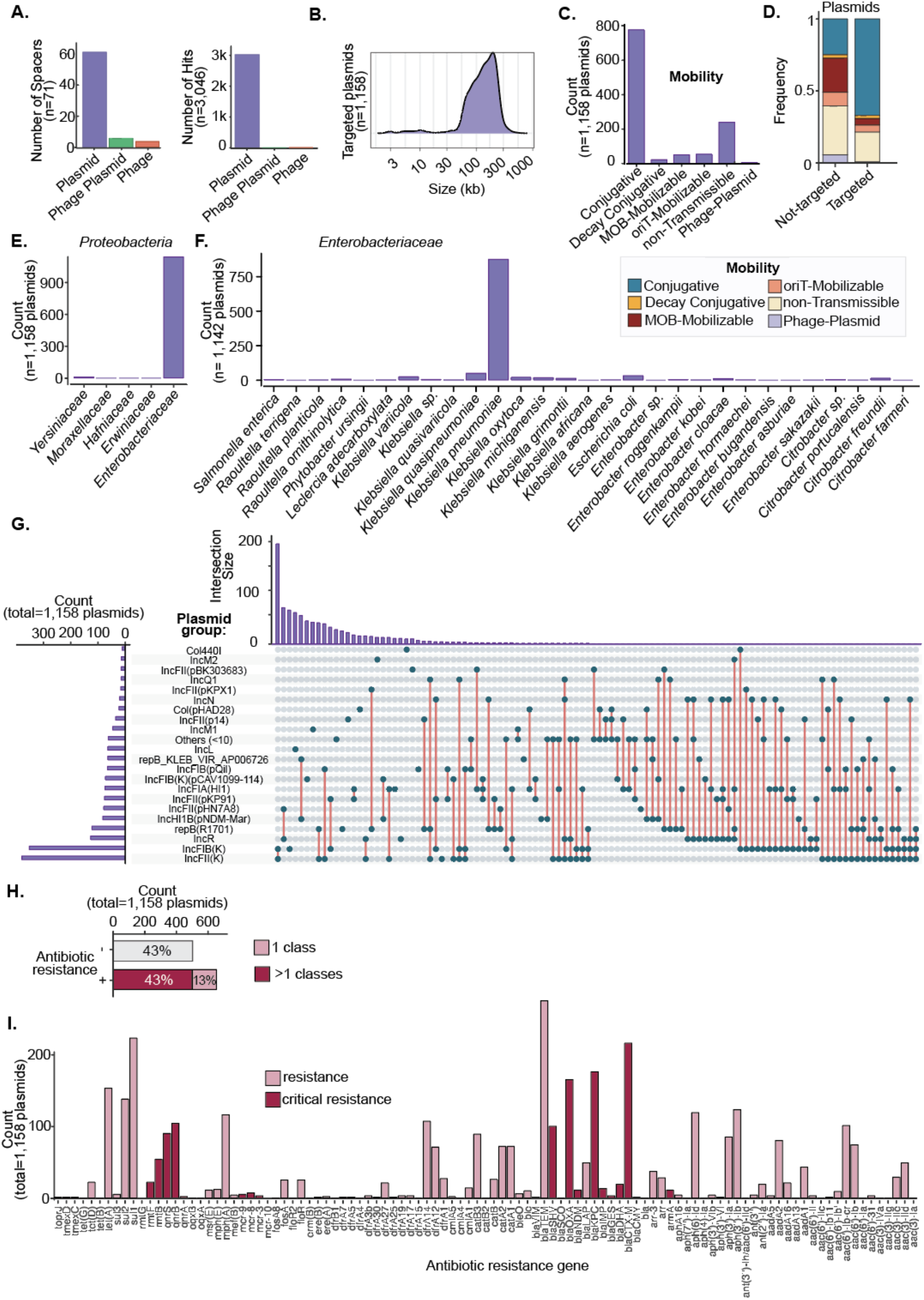
Characteristics of plasmids targeted by type IV-A3. **A)** Predicted targets of type IV-A3 systems based on spacer-protospacer matching analyses. The bars represent the counts of spacers matching sequences derived from plasmids, phage-plasmids, and phages. **B)** Size distribution of all targeted plasmids from the Refseq database. **C)** Mobility prediction of plasmids targeted by type IV-A3 spacers and non-targeted plasmids. **D)** Comparison of the predicted mobility type distributions of the RefSeq plasmids that are targeted and not targeted (non-targeted). Data under “targeted” is the same as in C. The mobility prediction of type IV-A3 carrying plasmids is shown as a reference. **E)** Family distribution of *Proteobacteria* hosts that harbor plasmids targeted by type IV-A3 spacers. **F)** Species distribution of *Enterobacteriaceae* hosts that harbor plasmids targeted by type IV-A3 spacers. **G)** Plasmid group distribution for plasmids targeted by type IV-A3 spacers. The upset plot shows the counts per group (left), and the intersection size of replicon combinations (right). Plasmid groups found to be targeted at low abundance (< 10 counts) have been grouped into the category “Others (<10 times)”. **H)** Antibiotic resistance carriage by plasmids targeted by type IV-A3 spacers. Counts of plasmids carrying more than five resistance genes are shown. The relative frequencies of each category are indicated inside the bars. **I)** Distribution of antibiotic resistance genes carried by plasmids encoding type IV-A3 CRISPR-Cas systems.

**Figure S7:**
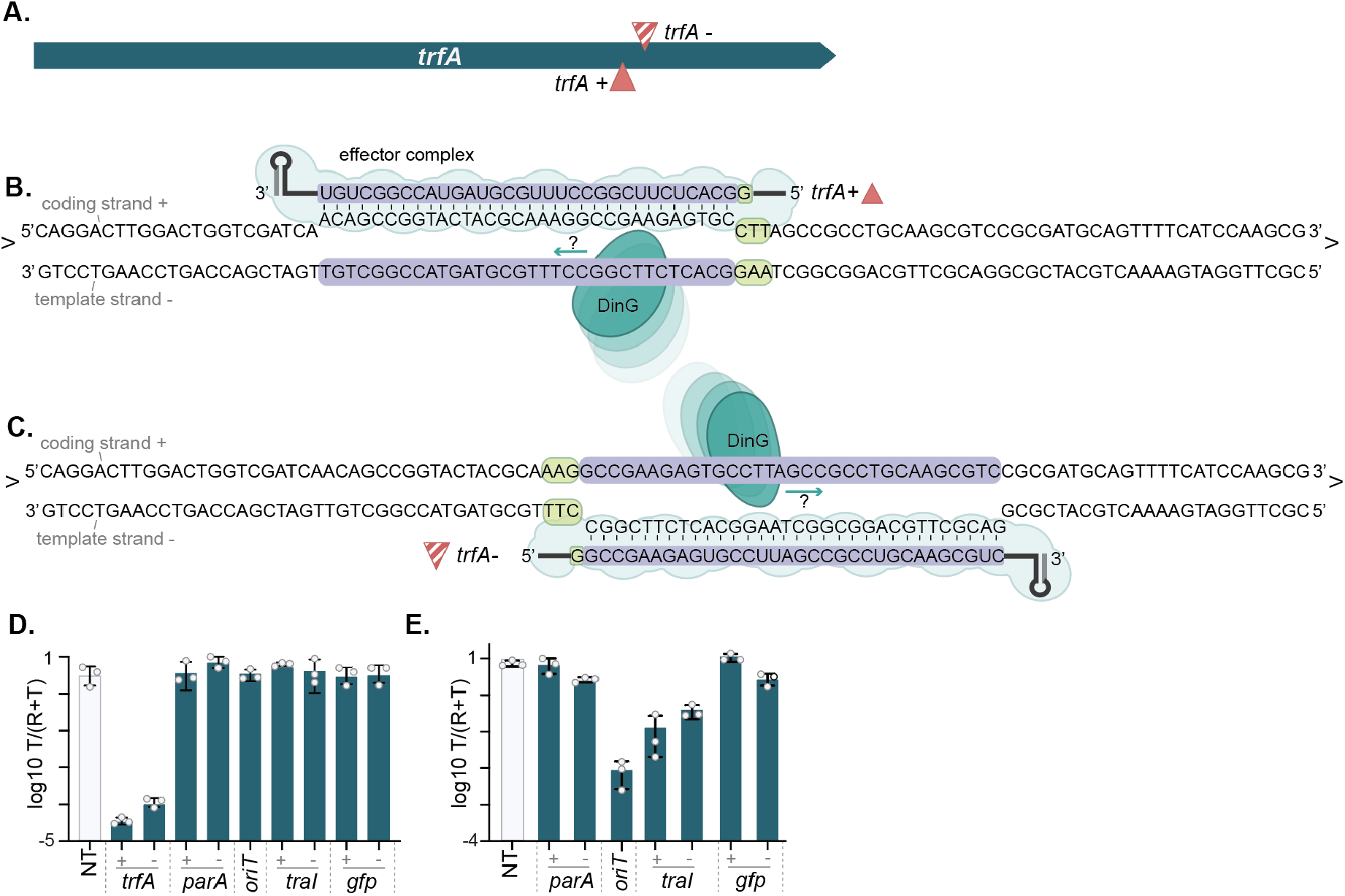
Targeting schematic and transconjugant frequencies under IV-A3 targeting. **A)** Schematic detailing the targeting of *trfA* with targeting crRNAs to exemplify the DNA strand nomenclature used throughout the targeting assays. Full triangles indicate crRNAs hybridizing to the coding strand (+) and crosshatched triangles to the template strand (-). **B)** Illustration of crRNA *trfA+*, exemplifying targeting of the coding strand, where the IV-A3 complex binds to the coding strand of the expressed gene. **C)** Illustration of crRNA *trfA-*, exemplifying targeting of the template strand, where the IV-A3 complex binds to the template strand of the expressed gene. In **B** and **C,** the protospacer and spacer sequences are indicated in purple, and the PAM sequence is indicated in green. **D)** Barplots representing the mean ratio of transconjugant (T) to recipient and transconjugant (R+T) in the incoming plasmid assay. **E)** Barplots representing the mean ratio of transconjugant (T) to recipient and transconjugant (R+T) in outgoing plasmid assay. In all experiments n = 3, error bars indicate SD.

**Figure S8:**
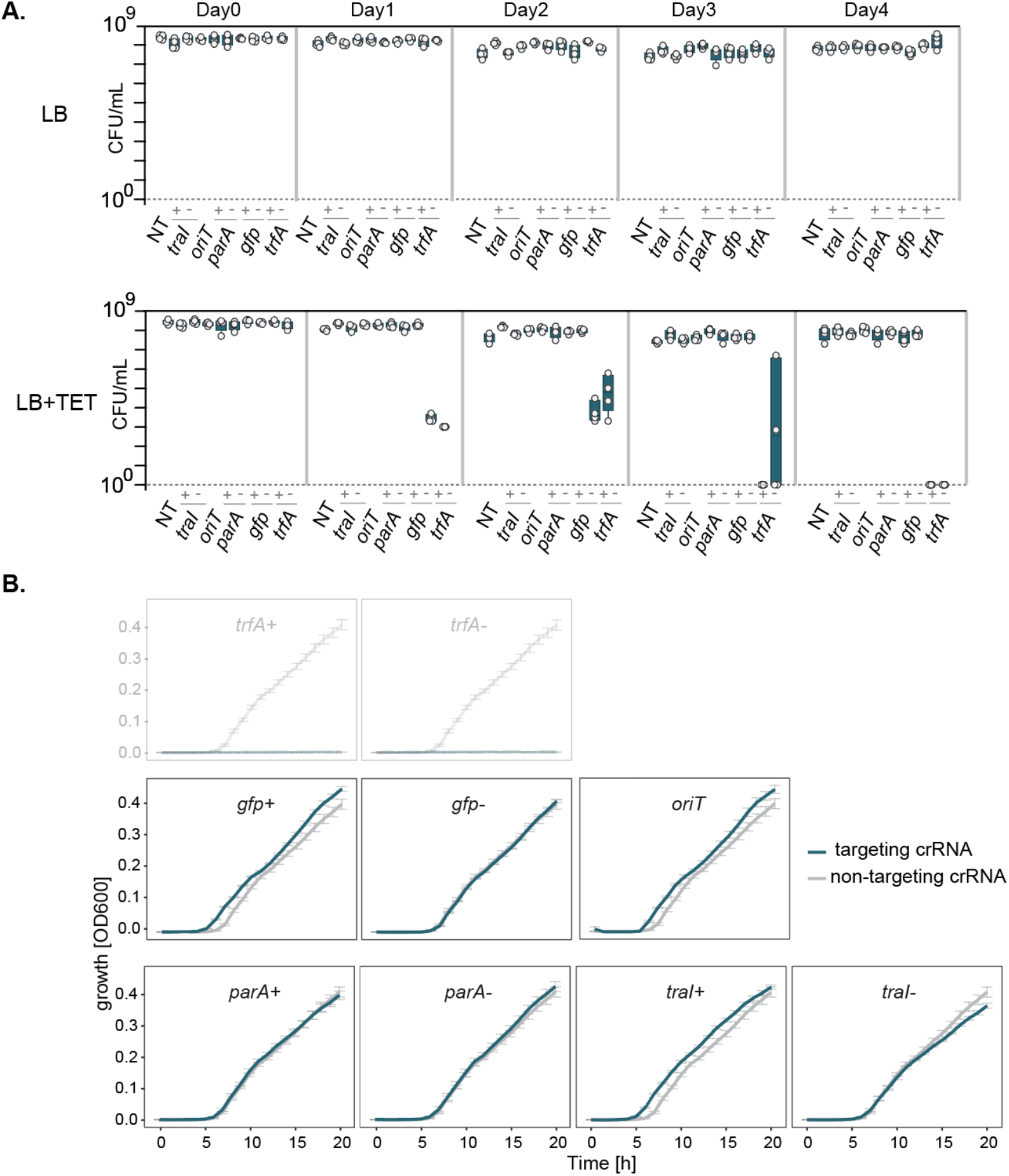
Plasmid stability and bacterial growth under IV-A3 targeting. **A)** pKJK5 stability under IV-A3 targeting and in the absence of antibiotic selection over four days (∼40 generations). Boxes represent the total CFU/ mL (plating on LB, top) or the CFU/ mL of pKJK5-carrying cells (selective plating on LB+TET, bottom) on each day (n=4); individual data points are indicated. **B)** Growth curves under pKJK5 targeting and antibiotic selection for pKJK5. Assessing the impact on growth during *trfA* targeting (top, grayed out) was unattainable due to the use of pKJK5 selection (tetracycline) and the instability of this plasmid under *trfA* targeting. Hourly OD_600_ measurements are indicated as a line representing the mean of five replicates, error bars indicate the SD. The NT control is included alongside each targeting crRNA to facilitate comparison.

**Figure S9.**
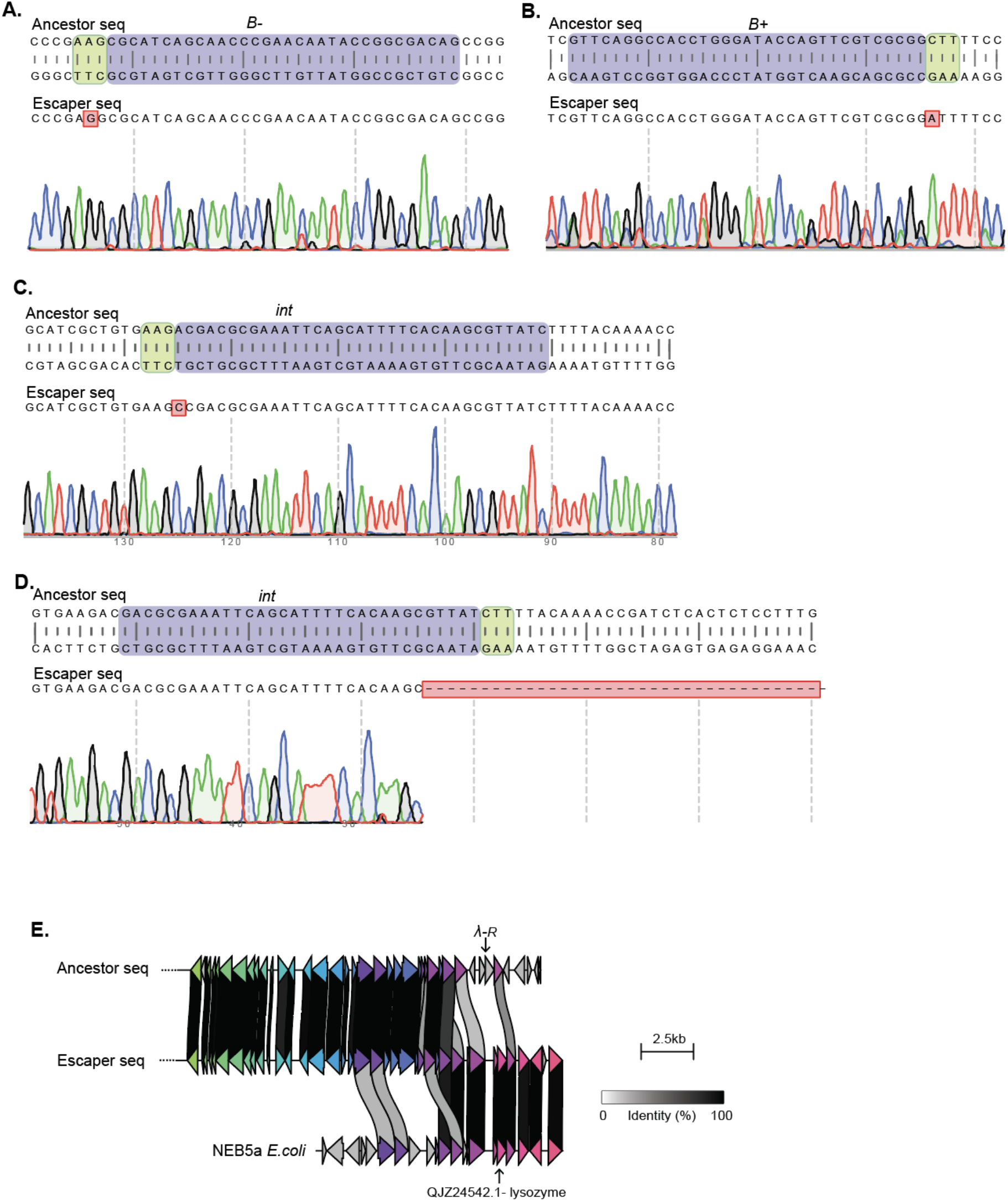
Phage-λ mutational evasion from IV-A3 interference. Ancestral phage sequences and the corresponding Sanger-sequencing results (below) for escaper phages evading gene B targeting, on the template **(A)** and the coding **(B)** strands, and two mutants from targeting either strand in the intergenic region **(C-D)**. **E)** Clinker alignment of selected genomic regions of the ancestral phage-λ, the isolated R-targeting escaper, and the homologous region present in other coliphages.

**Figure S10:**
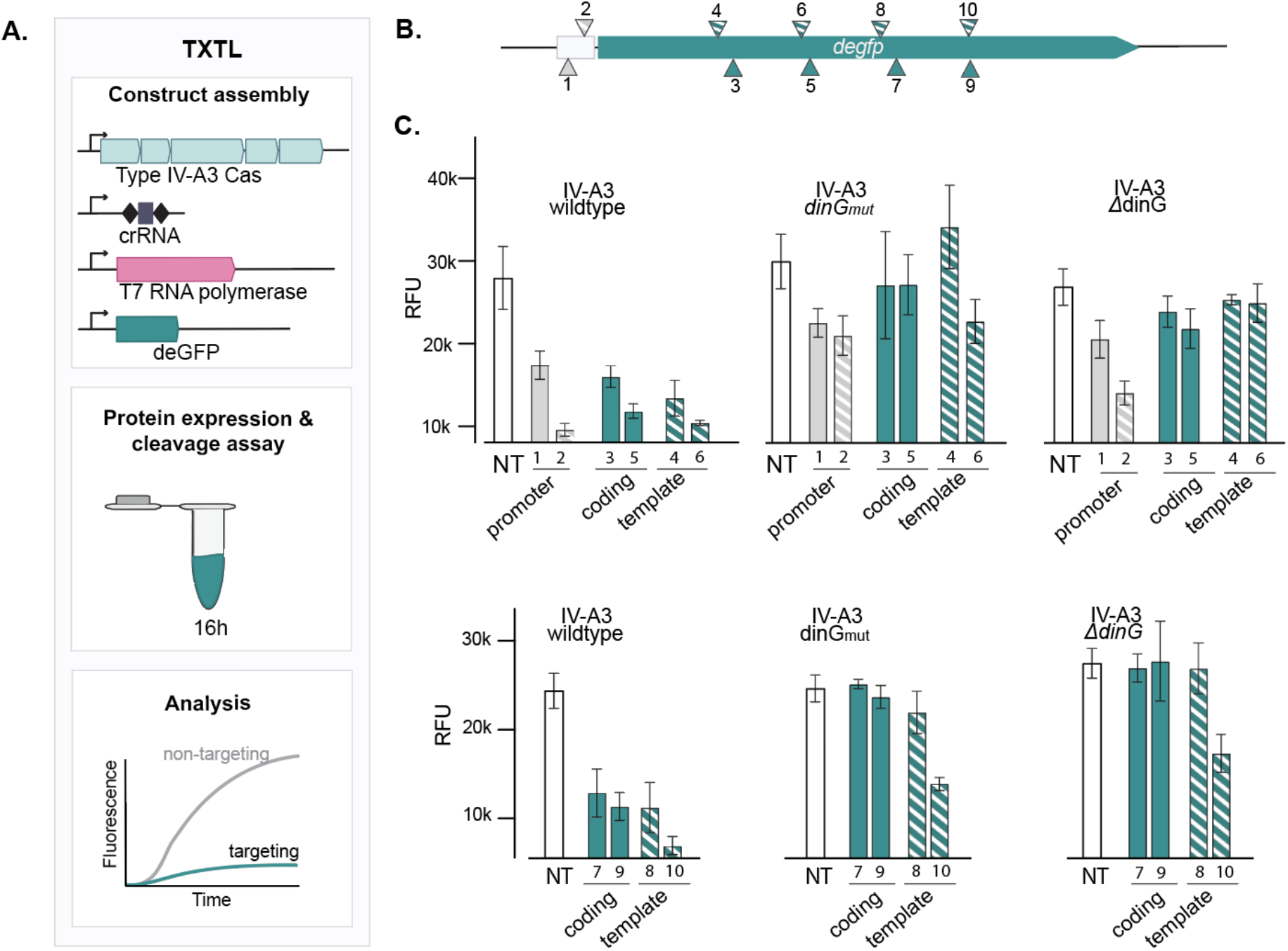
Assessment of type IV-A3 interference using an *in vitro* TXTL-based deGFP fluorescent reporter assay. **A)** Schematic of the TXTL experimental workflow: Plasmids encode the IV-A3 *cas* as well as one of the crRNAs (targeting *degfp* or control), the T7 RNA polymerase and *degfp*. CRISPR-Cas components are pre-expressed in TXTL prior to adding and expressing the targeted *degfp.* The rate of binding and the efficiency of transcriptional blocking impact the accumulation of deGFP and the resulting fluorescence of the TXTL reaction. **B)** Illustration of the approximate location where full triangles indicate crRNA hybridizing to the coding and crosshatched triangles to the template strand, respectively, for the promoter (gray) and *degfp* (green). **C)** Type IV-A3-mediated repression of deGFP production as RFU, performed in two independent experimental blocks (top and bottom). Multiple locations within the *degfp* gene and promoter regions were targeted, on both the coding (full) and template (crosshatched) strands. Bars show the mean of the relative fluorescence units (RFU) after 16 h (n=4) and error bars represent the SD.

**Figure S11:**
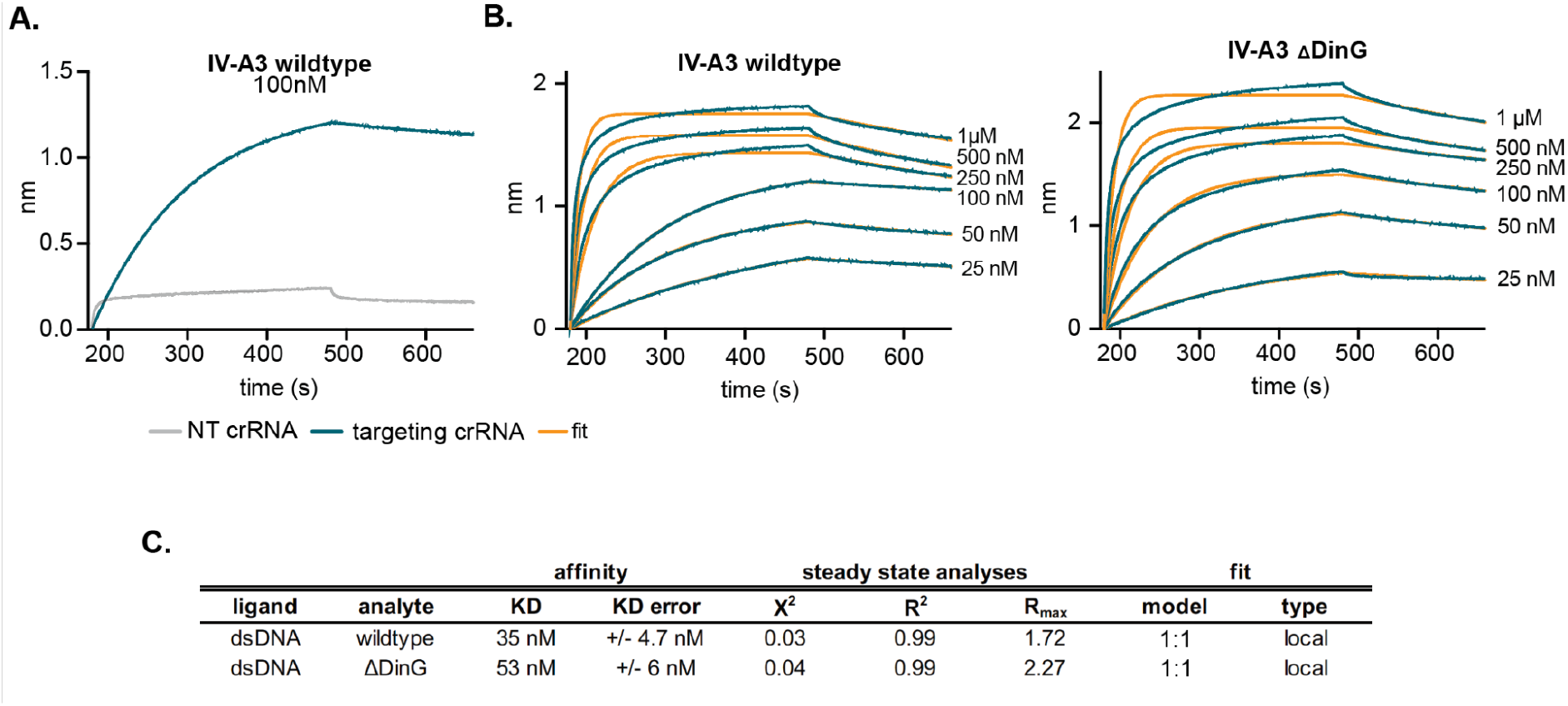
IV-A3 CRISPR-Cas complex affinity to double-stranded DNA targets. **A)** Bio-layer Interferometry (BLI) experimental data for the wildtype IV-A3 complex carrying a targeting (blue) or non-targeting (gray) crRNA guide. **B)** BLI experimental data (blue) and respective fittings (yellow) of the five titration steps for the wildtype (left) or ΔDinG (right) complex. Only association and dissociation steps are shown. **C)** Result table containing the steady-state analyses and fitting parameters.

**Figure S12:**
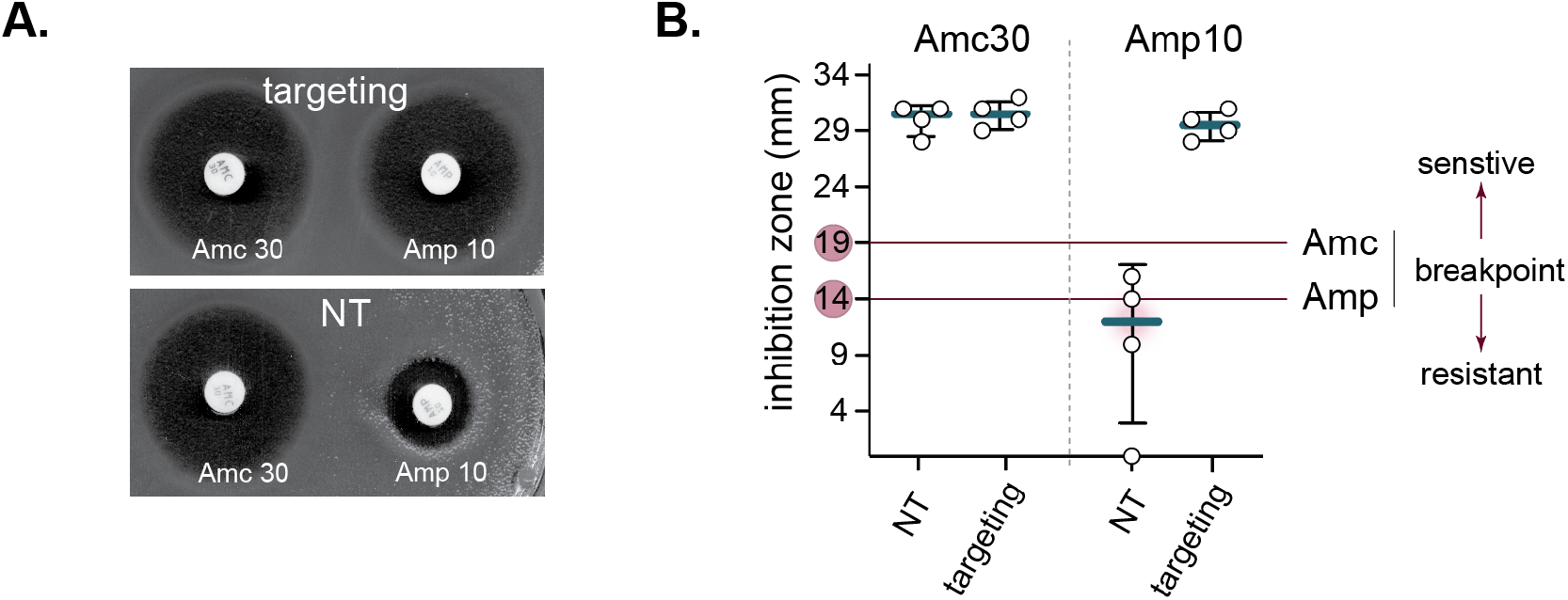
Experimental assessment of the re-sensitizing capacity of type IV-A3 CRISPR-Cas. **A)** Example picture of the disc diffusion MIC assay with (top, targeting) and without (bottom, NT) IV-A3 targeting of the chromosomally-encoded *bla*_SHV-187_ in *K. pneumoniae* 808330. In both treatments, we tested Amc (left, 30 µg; 20 µg amoxicillin + 10 µg β-lactamase inhibitor clavulanic acid) and Amp (right, 10 µg). **B)** Inhibition zones of disc diffusion MIC assay (exemplified in B). The EUCAST breakpoints for Amc and Amp are indicated in red.

**Figure S13:**
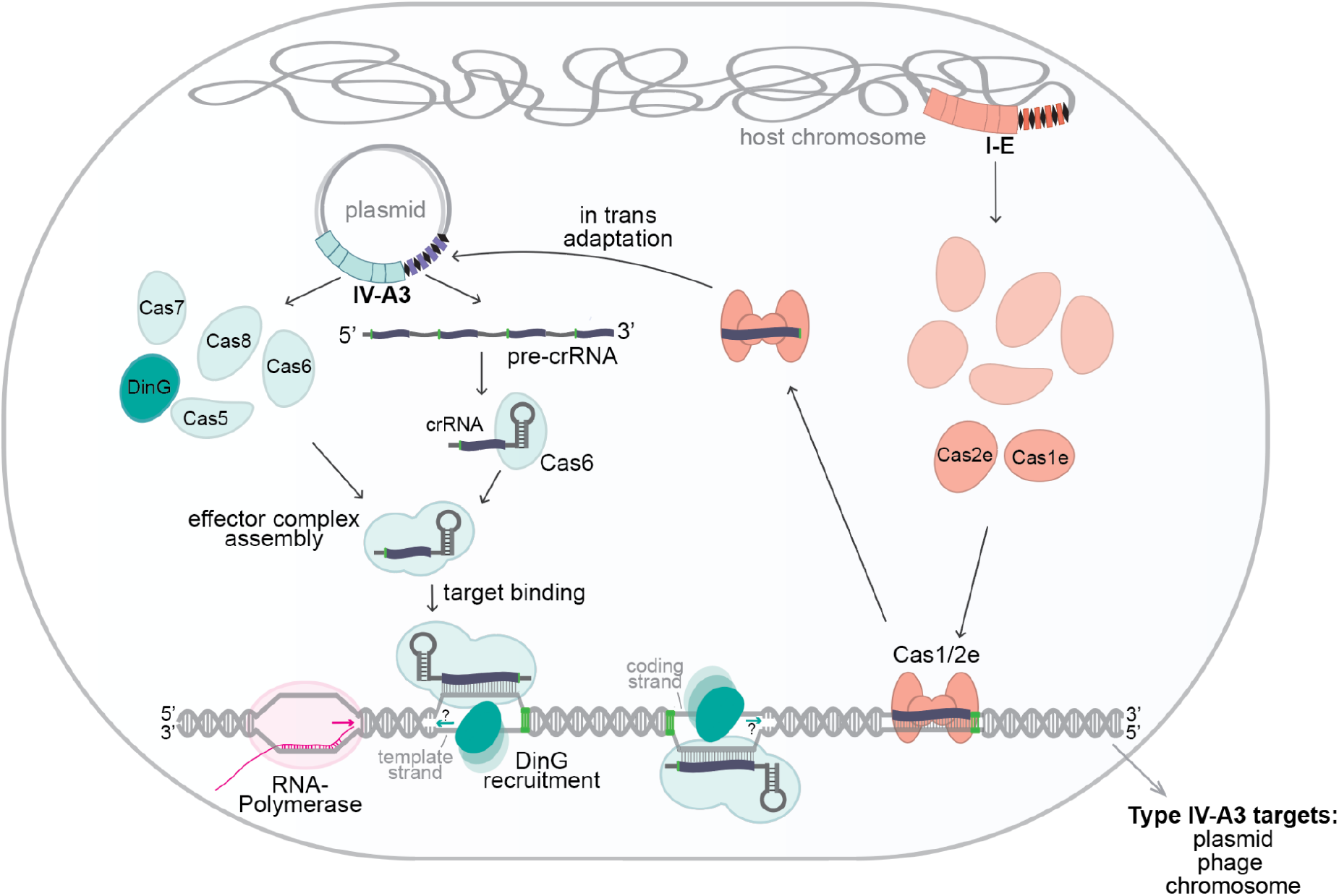
Proposed mechanistic model for spacer adaptation and target interference in type IV-A CRISPR-Cas systems. Spacer acquisition is facilitated by the co-option of the CRISPR-Cas I-E adaptation module (Cas1/Cas2e) that is encoded in the host chromosome. Type IV-A CRISPR and Cas components are expressed, and the pre-crRNA is subsequently processed by Cas6. The type IV-A effector complex assembles along the matured crRNAs and identifies protospacer targets in a PAM-dependent manner, presumably leading to DinG recruitment. Interference is elicited through transcriptional repression.

## REFERENCES

1. Shah, S. A., Erdmann, S., Mojica, F. J. M. & Garrett, R. A. Protospacer recognition motifs: mixed identities and functional diversity. RNA Biol. 10, 891–899 (2013).

2. Marraffini, L. A. & Sontheimer, E. J. Self versus non-self discrimination during CRISPR RNA-directed immunity. Nature 463, 568–571 (2010).

3. Makarova, K. S. et al. Evolutionary classification of CRISPR-Cas systems: a burst of class 2 and derived variants. Nat. Rev. Microbiol. 18, 67–83 (2020).

4. Liu, T. Y. & Doudna, J. A. Chemistry of Class 1 CRISPR-Cas effectors: Binding, editing, and regulation. J. Biol. Chem. 295, (2020).

5. Özcan, A. et al. Type IV CRISPR RNA processing and effector complex formation in Aromatoleum aromaticum. Nat Microbiol 4, 89–96 (2019).

6. Zhou, Y. et al. Structure of a type IV CRISPR-Cas ribonucleoprotein complex. iScience 24, 102201 (2021).

7. Pinilla-Redondo, R. et al. Type IV CRISPR-Cas systems are highly diverse and involved in competition between plasmids. Nucleic Acids Res. 48, 2000–2012 (2020).

8. Taylor, H. N. et al. Positioning Diverse Type IV Structures and Functions Within Class 1 CRISPR-Cas Systems. Front. Microbiol. 12, 671522 (2021).

9. Moya-Beltrán, A. et al. Evolution of Type IV CRISPR-Cas Systems: Insights from CRISPR Loci in Integrative Conjugative Elements of *Acidithiobacillia*. The CRISPR Journal vol. 4 656–672 Preprint at https://doi.org/10.1089/crispr.2021.0051 (2021).

10. Crowley, V. M. et al. A Type IV-A CRISPR-Cas System in *Pseudomonas aeruginosa* Mediates RNA-Guided Plasmid Interference *In Vivo*. The CRISPR Journal vol. 2 434–440 Preprint at https://doi.org/10.1089/crispr.2019.0048 (2019).

11. Newire, E., Aydin, A., Juma, S., Enne, V. I. & Roberts, A. P. Identification of a Type IV-A CRISPR-Cas System Located Exclusively on Plasmids in. Front. Microbiol. 11, 1937 (2020).

12. Guo, X. et al. Characterization of the self-targeting Type IV CRISPR interference system in Pseudomonas oleovorans. Nat Microbiol 7, 1870–1878 (2022).

13. Kamruzzaman, M. & Iredell, J. R. CRISPR-Cas System in Antibiotic Resistance Plasmids in. Front. Microbiol. 10, 2934 (2019).

14. Pitout, J. D. D. & Laupland, K. B. Extended-spectrum β-lactamase-producing Enterobacteriaceae: an emerging public-health concern. The Lancet Infectious Diseases vol. 8 159–166 Preprint at https://doi.org/10.1016/s1473-3099(08)70041-0 (2008).

15. Novais, A. et al. Evolutionary trajectories of beta-lactamase CTX-M-1 cluster enzymes: predicting antibiotic resistance. PLoS Pathog. 6, e1000735 (2010).

16. Shipman, S. L., Nivala, J., Macklis, J. D. & Church, G. M. Molecular recordings by directed CRISPR spacer acquisition. Science 353, aaf1175 (2016).

17. Swarts, D. C., Mosterd, C., van Passel, M. W. J. & Brouns, S. J. J. CRISPR Interference Directs Strand Specific Spacer Acquisition. PLoS ONE vol. 7 e35888 Preprint at https://doi.org/10.1371/journal.pone.0035888 (2012).

18. Yosef, I., Goren, M. G. & Qimron, U. Proteins and DNA elements essential for the CRISPR adaptation process in Escherichia coli. Nucleic Acids Research vol. 40 5569–5576 Preprint at https://doi.org/10.1093/nar/gks216 (2012).

19. Wang, J. et al. Structural and Mechanistic Basis of PAM-Dependent Spacer Acquisition in CRISPR-Cas Systems. Cell vol. 163 840–853 Preprint at https://doi.org/10.1016/j.cell.2015.10.008 (2015).

20. McKenzie, R. E., Almendros, C., Vink, J. N. A. & Brouns, S. J. J. Using CAPTURE to detect spacer acquisition in native CRISPR arrays. Nat. Protoc. 14, 976–990 (2019).

21. Díez-Villaseñor, C., Guzmán, N. M., Almendros, C., García-Martínez, J. & Mojica, F. J. M. CRISPR-spacer integration reporter plasmids reveal distinct genuine acquisition specificities among CRISPR-Cas I-E variants of Escherichia coli. RNA Biol. 10, 792–802 (2013).

22. Pinilla-Redondo, R. et al. CRISPR-Cas systems are widespread accessory elements across bacterial and archaeal plasmids. Nucleic Acids Research vol. 50 4315–4328 Preprint at https://doi.org/10.1093/nar/gkab859 (2022).

23. Bahl, M. I., Hansen, L. H., Licht, T. R. & Sørensen, S. J. Conjugative Transfer Facilitates Stable Maintenance of IncP-1 Plasmid pKJK5 in *Escherichia coli* Cells Colonizing the Gastrointestinal Tract of the Germfree Rat. Applied and Environmental Microbiology vol. 73 341–343 Preprint at https://doi.org/10.1128/aem.01971-06 (2007).

24. Klümper, U. et al. Broad host range plasmids can invade an unexpectedly diverse fraction of a soil bacterial community. The ISME Journal vol. 9 934–945 Preprint at https://doi.org/10.1038/ismej.2014.191 (2015).

25. Jaskólska, M., Adams, D. W. & Blokesch, M. Two defence systems eliminate plasmids from seventh pandemic Vibrio cholerae. Nature 604, 323–329 (2022).

26. Koopal, B. et al. Short prokaryotic Argonaute systems trigger cell death upon detection of invading DNA. Cell 185, 1471–1486.e19 (2022).

27. Deveau, H. et al. Phage response to CRISPR-encoded resistance in Streptococcus thermophilus. J. Bacteriol. 190, 1390–1400 (2008).

28. Sapranauskas, R. et al. The Streptococcus thermophilus CRISPR/Cas system provides immunity in Escherichia coli. Nucleic Acids Res. 39, 9275–9282 (2011).

29. Semenova, E. et al. Interference by clustered regularly interspaced short palindromic repeat (CRISPR) RNA is governed by a seed sequence. Proc. Natl. Acad. Sci. U. S. A. 108, 10098–10103 (2011).

30. Wimmer, F., Mougiakos, I., Englert, F. & Beisel, C. L. Rapid cell-free characterization of multi-subunit CRISPR effectors and transposons. Mol. Cell 82, 1210–1224.e6 (2022).

31. Antimicrobial Resistance Collaborators. Global burden of bacterial antimicrobial resistance in 2019: a systematic analysis. Lancet 399, 629–655 (2022).

32. Edgar, R., Friedman, N., Molshanski-Mor, S. & Qimron, U. Reversing bacterial resistance to antibiotics by phage-mediated delivery of dominant sensitive genes. Appl. Environ. Microbiol. 78, 744–751 (2012).

33. Hochvaldová, L. et al. Restoration of antibacterial activity of inactive antibiotics via combined treatment with a cyanographene/Ag nanohybrid. Sci. Rep. 12, 5222 (2022).

34. Bikard, D. & Barrangou, R. Using CRISPR-Cas systems as antimicrobials. Curr. Opin. Microbiol. 37, 155–160 (2017).

35. Benz, F. & Hall, A. R. Host-specific plasmid evolution explains the variable spread of clinical antibiotic-resistance plasmids. Proc. Natl. Acad. Sci. U. S. A. 120, e2212147120 (2023).

36. Benz, F. et al. Plasmid- and strain-specific factors drive variation in ESBL-plasmid spread in vitro and in vivo. ISME J. 15, 862–878 (2021).

37. Bernheim, A., Bikard, D., Touchon, M. & Rocha, E. P. C. Atypical organizations and epistatic interactions of CRISPRs and cas clusters in genomes and their mobile genetic elements. Nucleic Acids Res. 48, 748–760 (2020).

38. Peters, J. E., Makarova, K. S., Shmakov, S. & Koonin, E. V. Recruitment of CRISPR-Cas systems by Tn7-like transposons. Proc. Natl. Acad. Sci. U. S. A. 114, E7358–E7366 (2017).

39. Al-Shayeb, B., et al. Diverse virus-encoded CRISPR-Cas systems include streamlined genome editors. Cell 185, 4574–4586.e16 (2022).

40. Weissman, J. L., Stoltzfus, A., Westra, E. R. & Johnson, P. L. F. Avoidance of Self during CRISPR Immunization. Trends in Microbiology vol. 28 543–553 Preprint at https://doi.org/10.1016/j.tim.2020.02.005 (2020).

41. Klompe, S. E., Vo, P. L. H., Halpin-Healy, T. S. & Sternberg, S. H. Transposon-encoded CRISPR-Cas systems direct RNA-guided DNA integration. Nature 571, 219–225 (2019).

42. Strecker, J. et al. RNA-guided DNA insertion with CRISPR-associated transposases. Science 365, 48–53 (2019).

43. Rybarski, J. R., Hu, K., Hill, A. M., Wilke, C. O. & Finkelstein, I. J. Metagenomic discovery of CRISPR-associated transposons. Proc. Natl. Acad. Sci. U. S. A. 118, (2021).

44. Al-Shayeb, B. et al. Clades of huge phages from across Earth’s ecosystems. Nature 578, 425–431 (2020).

45. Wu, W. Y. et al. The miniature CRISPR-Cas12m effector binds DNA to block transcription. Mol. Cell 82, 4487–4502.e7 (2022).

46. Tesson, F. & Bernheim, A. Synergy and regulation of antiphage systems: toward the existence of a bacterial immune system? Curr. Opin. Microbiol. 71, 102238 (2023).

47. Rocha, E. P. C. & Bikard, D. Microbial defenses against mobile genetic elements and viruses: Who defends whom from what? PLoS Biol. 20, e3001514 (2022).

48. Little, J. W. & Mount, D. W. The SOS regulatory system of Escherichia coli. Cell 29, 11–22 (1982).

49. Janion, C. Some aspects of the SOS response system--a critical survey. Acta Biochim. Pol. 48, 599–610 (2001).

50. Malone, L. M., Hampton, H. G., Morgan, X. C. & Fineran, P. C. Type I CRISPR-Cas provides robust immunity but incomplete attenuation of phage-induced cellular stress. Nucleic Acids Res. 50, 160–174 (2022).

51. Wegrzyn, G. & Wegrzyn, A. Stress responses and replication of plasmids in bacterial cells. Microb. Cell Fact. 1, 2 (2002).

52. Huang, C. J., Adler, B. A. & Doudna, J. A. A naturally DNase-free CRISPR-Cas12c enzyme silences gene expression. Mol. Cell 82, 2148–2160.e4 (2022).

53. Quinones-Olvera, N., et al. Diverse and abundant viruses exploit conjugative plasmids. bioRxiv (2023) doi:10.1101/2023.03.19.532758.

54. Qi, L. S. et al. Repurposing CRISPR as an RNA-guided platform for sequence-specific control of gene expression. Cell 152, 1173–1183 (2013).

55. Bikard, D. et al. Programmable repression and activation of bacterial gene expression using an engineered CRISPR-Cas system. Nucleic Acids Res. 41, 7429–7437 (2013).

56. Kim, S. K. et al. Efficient Transcriptional Gene Repression by Type V-A CRISPR-Cpf1 from Eubacterium eligens. ACS Synth. Biol. 6, 1273–1282 (2017).

57. Zhang, X. et al. Multiplex gene regulation by CRISPR-ddCpf1. Cell Discov 3, 17018 (2017).

58. Luo, M. L., Mullis, A. S., Leenay, R. T. & Beisel, C. L. Repurposing endogenous type I CRISPR-Cas systems for programmable gene repression. Nucleic Acids Res. 43, 674–681 (2015).

59. Cassini, A. et al. Attributable deaths and disability-adjusted life-years caused by infections with antibiotic-resistant bacteria in the EU and the European Economic Area in 2015: a population-level modelling analysis. Lancet Infect. Dis. 19, 56–66 (2019).

60. Bikard, D. et al. Exploiting CRISPR-Cas nucleases to produce sequence-specific antimicrobials. Nat. Biotechnol. 32, 1146–1150 (2014).

61. Uribe, R. V. et al. Bacterial resistance to CRISPR-Cas antimicrobials. Sci. Rep. 11, 17267 (2021).

62. Kroll, J., Klinter, S., Schneider, C., Voss, I. & Steinbüchel, A. Plasmid addiction systems: perspectives and applications in biotechnology. Microb. Biotechnol. 3, 634–657 (2010).

63. Citorik, R. J., Mimee, M. & Lu, T. K. Sequence-specific antimicrobials using efficiently delivered RNA-guided nucleases. Nat. Biotechnol. 32, 1141–1145 (2014).

64. Wick, R. R., Judd, L. M., Gorrie, C. L. & Holt, K. E. Unicycler: Resolving bacterial genome assemblies from short and long sequencing reads. PLoS Comput. Biol. 13, e1005595 (2017).

65. Russel, J., Pinilla-Redondo, R., Mayo-Muñoz, D., Shah, S. A. & Sørensen, S. J. CRISPRCasTyper: Automated Identification, Annotation, and Classification of CRISPR-Cas Loci. CRISPR J 3, 462–469 (2020).

66. Carattoli, A. et al. In silico detection and typing of plasmids using PlasmidFinder and plasmid multilocus sequence typing. Antimicrob. Agents Chemother. 58, (2014).

67. Robertson, J. & Nash, J. H. E. MOB-suite: software tools for clustering, reconstruction and typing of plasmids from draft assemblies. Microbial Genomics 4, e000206 (2018).

68. Alcock, B. P. et al. CARD 2023: expanded curation, support for machine learning, and resistome prediction at the Comprehensive Antibiotic Resistance Database. Nucleic Acids Res. 51, (2023).

69. Huisman, J. S. et al. Estimating plasmid conjugation rates: A new computational tool and a critical comparison of methods. Plasmid 121, 102627 (2022).

70. Katoh, K. & Standley, D. M. MAFFT multiple sequence alignment software version 7: improvements in performance and usability. Mol. Biol. Evol. 30, 772–780 (2013).

71. Price, M. N., Dehal, P. S. & Arkin, A. P. FastTree 2--approximately maximum-likelihood trees for large alignments. PLoS One 5, e9490 (2010).

72. Letunic, I. & Bork, P. Interactive Tree Of Life (iTOL) v5: an online tool for phylogenetic tree display and annotation. Nucleic Acids Res. 49, W293–W296 (2021).

73. Blackwell, G. A. et al. Exploring bacterial diversity via a curated and searchable snapshot of archived DNA sequences. PLoS Biol. 19, e3001421 (2021).

74. Li, W. & Godzik, A. Cd-hit: a fast program for clustering and comparing large sets of protein or nucleotide sequences. Bioinformatics 22, 1658–1659 (2006).

75. Camacho C., Coulouris G., Avagyan V., Ma N., Papadopoulos J., Bealer K., Madden T.L. BLAST+: architecture and applications. BMC Bioinformatics 10,.

76. Schmartz, G. P. et al. PLSDB: advancing a comprehensive database of bacterial plasmids. Nucleic Acids Res. 50, D273–D278 (2022).

77. Feldgarden, M. et al. AMRFinderPlus and the Reference Gene Catalog facilitate examination of the genomic links among antimicrobial resistance, stress response, and virulence. Sci. Rep. 11, 12728 (2021).

78. Pfeifer, E., Bonnin, R. A. & Rocha, E. P. C. Phage-Plasmids Spread Antibiotic Resistance Genes through Infection and Lysogenic Conversion. MBio 13, e0185122 (2022).

79. Cury, J., Abby, S. S., Doppelt-Azeroual, O., Néron, B. & Rocha, E. P. C. Identifying Conjugative Plasmids and Integrative Conjugative Elements with CONJscan. Methods Mol. Biol. 2075, 265–283 (2020).

80. Ares-Arroyo, M., Coluzzi, C. & Rocha, E. P. C. Origins of transfer establish networks of functional dependencies for plasmid transfer by conjugation. Nucleic Acids Res. 51, 3001–3016 (2023).

81. Wickham, H. ggplot2. (Springer International Publishing).

82. Conway, J. R., Lex, A. & Gehlenborg, N. UpSetR: an R package for the visualization of intersecting sets and their properties. Bioinformatics 33, 2938–2940 (2017).

83. Wilke, C. ggridges: Ridgeline Plots in ‘ggplot2’. R package version 0.5.4,. https://wilkelab.org/ggridges/. (2022).

84. Martin, M. Cutadapt removes adapter sequences from high-throughput sequencing reads. EMBnet.journal 17, 10–12 (2011).

85. Kim, D., Langmead, B. & Salzberg, S. L. HISAT: a fast spliced aligner with low memory requirements. Nat. Methods 12, 357–360 (2015).

86. Thrash, A., Arick, M. & Peterson, D. G. Quack: A quality assurance tool for high throughput sequence data. Anal. Biochem. 548, (2018).

87. Mortazavi, A., Williams, B. A., McCue, K., Schaeffer, L. & Wold, B. Mapping and quantifying mammalian transcriptomes by RNA-Seq. Nat. Methods 5, (2008).

88. Wickham, H. ggplot2: Elegant Graphics for Data Analysis. (Springer Science & Business Media, 2009).

89. Datsenko, K. A. & Wanner, B. L. One-step inactivation of chromosomal genes in Escherichia coli K-12 using PCR products. Proc. Natl. Acad. Sci. U. S. A. 97, 6640–6645 (2000).

90. Bolger, A. M., Lohse, M. & Usadel, B. Trimmomatic: a flexible trimmer for Illumina sequence data. Bioinformatics 30, 2114–2120 (2014).

91. Zhang, J., Kobert, K., Flouri, T. & Stamatakis, A. PEAR: a fast and accurate Illumina Paired-End reAd mergeR. Bioinformatics 30, 614–620 (2014).

92. Jakočiūnė, D. & Moodley, A. A Rapid Bacteriophage DNA Extraction Method. Methods Protoc 1, (2018).

93. Brettin, T. et al. RASTtk: a modular and extensible implementation of the RAST algorithm for building custom annotation pipelines and annotating batches of genomes. Sci. Rep. 5, 8365 (2015).

94. Gilchrist, C. L. M. & Chooi, Y.-H. clinker & clustermap.js: automatic generation of gene cluster comparison figures. Bioinformatics 37, 2473–2475 (2021).

95. Leenay, R. T. et al. Identifying and Visualizing Functional PAM Diversity across CRISPR-Cas Systems. Mol. Cell 62, 137–147 (2016).

96. Ondov, B. D., Bergman, N. H. & Phillippy, A. M. Interactive metagenomic visualization in a Web browser. BMC Bioinformatics 12, 1–10 (2011).

97. Wiegand, I., Hilpert, K. & Hancock, R. E. Agar and broth dilution methods to determine the minimal inhibitory concentration (MIC) of antimicrobial substances. Nat. Protoc. 3, (2008).

98. ESCMID-European Society of Clinical Microbiology & Diseases, I. eucast: MIC determination. https://www.eucast.org/ast_of_bacteria/mic_determination.

99. ESCMID-European Society of Clinical Microbiology & Diseases, I. eucast: Clinical breakpoints and dosing of antibiotics. https://www.eucast.org/clinical_breakpoints.

100. Morris, D. et al. Inter-hospital outbreak of Klebsiella pneumoniae producing KPC-2 carbapenemase in Ireland. J. Antimicrob. Chemother. 67, 2367–2372 (2012).

101. Diard, M. et al. Inflammation boosts bacteriophage transfer between spp. Science 355, 1211–1215 (2017).

